# Keap1 binds cytokine promoters upon virus infection and moderates their induction by recruiting NFκB p50 and G9a-GLP

**DOI:** 10.1101/2020.09.30.321703

**Authors:** Veronica Elizabeth Burns, Tom Klaus Kerppola

**Affiliations:** Department of Biological Chemistry, University of Michigan, Ann Arbor, MI 48109-5606

## Abstract

Innate immunity requires a balance of positive and negative regulators of cytokine transcription. Keap1 deletion in mice alters innate immunity and inflammation. We investigated the influence of Keap1 on cytokine gene induction by Sendai virus infection in mouse embryo fibroblasts (MEFs). Keap1 bound to the *Ifnb1, Tnf* and *Il6* promoters upon virus infection, and moderated viral induction of their transcription. Keap1 was required for viral induction of NFκB p50 and G9a-GLP lysine methyltransferase binding to these genes. Keap1 formed BiFC complexes with NFκB p50 that were localized to the nuclei in a subset of cells. Nrf2 counteracted viral induction of Keap1 binding to the promoters, and the effects of Keap1 on NFκB p50 and on G9a-GLP recruitment. Lysine methyltransferase inhibitors enhanced viral induction of transcription of the genes that were bound by Keap1 only in MEFs with intact Keap1, and not in *Keap1-/-* MEFs. They also enhanced NFκB p50 and NFκB p65 recruitment to these genes only in MEFs with intact Keap1, whereas they inhibited G9a-GLP recruitment. The reciprocal effects of Keap1 and of G9a-GLP lysine methyltransferase activity on chromatin binding by each other constitute a feedback circuit that moderates viral induction of cytokine transcription.

**Summary:** Virus infection induces Keap1 binding to cytokine promoters, which recruits NFκB p50 and G9a-GLP and moderates their transcription.

## Introduction

Innate immune and inflammatory responses are coordinated by cytokines. Many mechanisms that that activate cytokine transcription have been described. Robust control mechanisms require a balance of positive and negative effectors. The mechanisms that moderate cytokine transcription are incompletely characterized (*1- 4*).

The effects of transcription factor mutations on innate immune response gene expression have been investigated in mouse embryo fibroblasts (MEFs) (*5, 6*). Studies in MEFs circumvent the indirect effects of changes in development and in immune sensitization. Sendai virus infection induces a broad range of innate immune response genes in fibroblasts by activating RIG-I and other pattern recognition receptors (*5, 7, 8*). Numerous genes that influence immune functions are activated in fibroblasts in different organs of virus infected mice (*9*). Many of these genes exhibit chromatin accessibility and histone modifications consistent with activation potential also in fibroblasts from uninfected mice. Fibroblasts participate in immune responses *in vivo* and are a valid model for investigation of the mechanisms for immune gene regulation *in vitro*.

The NFκB p50 and p65 proteins form dimers that can regulate many immunomodulatory genes. Mutations in the genes that encode NFκB p50 *versus* NFκB p65 have distinct effects on cytokine induction in MEFs (*5, 6*). NFκB p50 can moderate cytokine expression in many cell types, and can prevent autoimmune and inflammatory responses in mice (*3, 10-13*). NFκB p50 and NFκB p65 bind and regulate overlapping sets of genes (*14*). Their relative levels of binding vary in response to various stimuli. NFκB p50 phosphorylation can regulate in vitro binding to oligonucleotides (*15-17*). The mechanisms that control the selective recruitment of NFκB p50 *versus* NFκB p65 to specific promoters in response to immunomodulatory signals are not known.

The low basal levels of cytokine transcription in MEFs compared to dendritic cells correlate with higher levels of histone H3 lysine 9 dimethylation (H3K9me2) in MEFs. Deletion of the G9a or GLP subunit of the H3 lysine methyltransferase enhances poly(I:C) or TNFα induction of interferon-responsive gene transcription (*3, 4*). NFκB p50 and RelB coprecipitate with GLP and G9a, respectively, and can influence their binding to cytokine genes, suggesting that they can moderate cytokine induction in concert (*3, 18*). The factors that control the corecruitment of G9a-GLP and NFκB subunits to cytokine promoters and the roles of lysine methyltransferase activity in their recruitment are not known.

Keap1 and its interaction partner Nrf2 regulate electrophile response genes (*19, 20*). They are also implicated in the regulation of innate immunity by the altered autoimmune and inflammatory responses in mice that lack Keap1 or Nrf2 (*21-26*). Keap1 as well as Nrf2 deficiency can cause both pro- and anti-inflammatory dysfunctions (*27-33*). The complex effects of Keap1 and Nrf2 deficiencies on inflammation suggest that they influence immune responses through many mechanisms. The effects of Keap1 deficiency on immune and inflammatory responses are often interpreted to be due to increased Nrf2 activity (*34-37*). This interpretation assumes that the effects of Keap1 on innate immune responses are related to the mechanism for Keap1 regulation of electrophile response gene transcription. The presumption that Keap1 effects must be mediated by Nrf2 is unwarranted in cases where the molecular mechanisms are unknown.

Keap1 depletion has contrasting effects on cytokine transcription under different conditions. Keap1 depletion increases cytokine transcription induced by LPS and by *Mycobacterium avium* infection in mouse macrophage and human monocyte cell lines, and in human primary macrophages, respectively (*27, 28*). In contrast, conditional *Keap1*^*fl/fl*^ deletion reduces cytokine transcription induced by LPS/IFN-γ in bone marrow derived macrophages (*36*). The contrasting effects of Keap1 depletion in different experiments indicate either that the indirect effects of Keap1 vary between these cells, or that Keap1 regulates cytokine transcription through additional, previously undescribed mechanisms.

Whereas Keap1 is enriched in the cytoplasm, Keap1 can interact with nuclear proteins and regulates their functions in DNA repair and replication (*38, 39*). Keap1 can interact with NFκB p65 in yeast and in mammalian cell extracts (*40*). Keap1 can also interact with NFκB p50 and NFκB p65 in liver extracts from rabbits infected with the Rabbit Hemorrhagic Disease Virus, and the distribution of Keap1 overlaps with that of NFκB p50 in liver cell nuclei (*41*). It is likely that Keap1 interacts with different partners in different subcellular locations, but the distributions of the complexes that Keap1 forms with specific partners have not been analyzed.

Drosophila Keap1 binds to specific genes on polytene chromosomes and regulates their transcription (*42, 43*). The observation that Drosophila Keap1 binds to many loci that contain innate immune response genes, suggested the possibility that Keap1 regulates immunomodulatory genes by binding to chromatin.

We found that Sendai virus infection induces Keap1 binding to selected cytokine promoters, which moderates their induction in MEFs. Keap1 is essential for NFκB p50 and for G9a-GLP lysine methyltransferase recruitment to cytokine genes. G9a-GLP inhibitors enhance viral induction of cytokine transcription and Keap1 and NFκB recruitment. Keap1 forms complexes with NFκB p50 and with NFκB p65 in living cells.

## Results

### Keap1 moderates viral induction of cytokine transcription

We investigated the roles of Keap1 in cytokine gene induction by Sendai virus infection. We focused on virus induction of the *Ifnb1, Tnf* and *Il6* genes in primary MEFs to circumvent indirect effects of changes in development and in immune sensitization. These genes represent different classes of cytokine regulatory mechanisms, as reflected by differences in the transcription factors, the chromatin compositions, and the functions of nucleosome remodeling at these promoters (*44, 45*). We tested the effects of *Keap1-/-* deletions in MEFs with *Nrf2-/-* deletions to distinguish the direct effects of Keap1 from indirect effects of Nrf2 activation (Fig. 1, S1).

**Fig 1.**
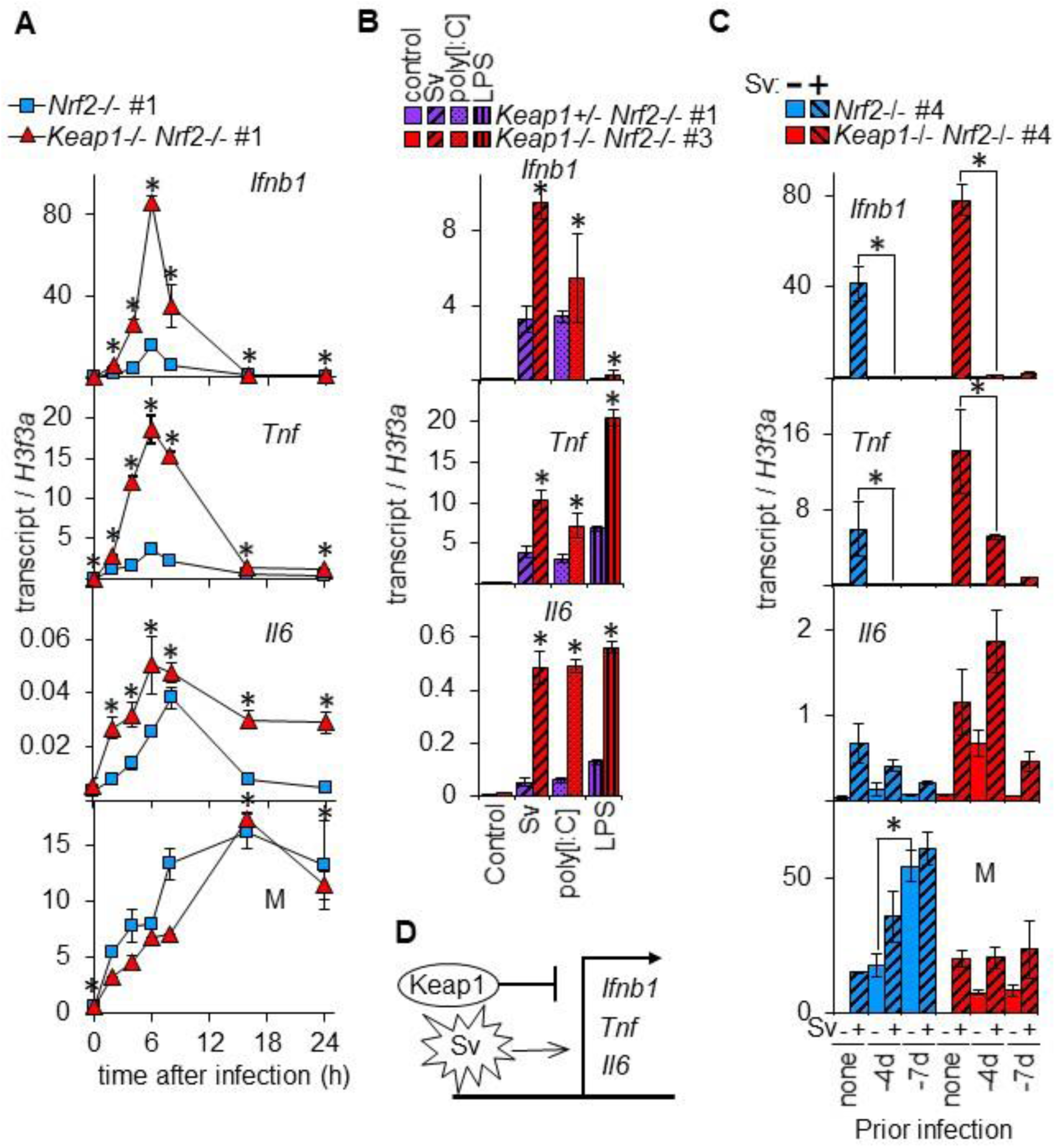
*Keap1-/-* deletions increase viral induction of *Ifnb1, Tnf* and *Il6* transcription independently of Nrf2. (**A**) The levels of *Ifnb1, Tnf, Il6*, and Sendai virus *M* gene transcripts were measured in *Nrf2-/- #1*, and in *Keap1-/- Nrf2-/- #1* MEFs at the indicated times after Sendai virus infection. The statistical significance of the differences in transcript levels were tested by two-factor ANOVA analyses of data from 7-10 experiments that were performed using 6-7 different sets of MEFs (* p<0.0001). MEFs from different embryos are identified by #number. Experiments with different MEFs, including *Keap1-/-* MEFs with intact Nrf2 are shown in Fig. S1A-S1C. (**B**) The levels of *Ifnb1, Tnf*, and *Il6* gene transcripts were measured in *Keap1+/- Nrf2-/-* #1 and in *Keap1-/- Nrf2-/-* #1 MEFs: 6 h after Sendai virus infection; 6 h after 4 μg/ml poly[I:C] addition; 2 h after 0.2 μl/ml lipopolysaccharide (LPS) addition; and in vehicle controls (con) as indicated below the bars. The statistical significance was tested by two-factor ANOVAs of data from 3 experiments with 2 different pairs of MEFs (* p<0.0001). (**C**) The levels of *Ifnb1, Tnf, Il6* and *M* gene transcripts were measured in *Nrf2-/-* #4, and in *Keap1-/- Nrf2-/-* #4 MEFs 6 h after Sendai virus infection following either no prior infection, or prior infections 4 days or 7 days earlier as indicates below the bars. The statistical significance was tested by two-factor ANOVAs of data from 2 experiments with different pairs of MEFs (* p<0.001). (**D**) Effects of Keap1 on *Ifnb1, Tnf* and *Il6* induction by Sendai virus (Sv) infection.

The levels of *Ifnb1, Tnf*, and *Il6* transcripts increased more rapidly and reached higher levels in *Keap1-/- Nrf2-/-* MEFs than in *Nrf2-/-* MEFs following Sendai virus infection (Fig. 1A). To evaluate the reproducibility of the effects of *Keap1-/-* deletions, we compared the *Ifnb1, Tnf* and *Il6* transcript levels in MEFs that were isolated from at least 5 embryos of each genotype. The peak *Ifnb1, Tnf*, and *Il6* transcript levels were 3 ± 0.5, 8 ± 3, and 3 ± 0.9 fold higher, respectively, in *Keap1-/- Nrf2-/-* MEFs than in *Nrf2-/-* MEFs (Fig. 1A, S1A-C, 5A, S5A; mean ± sem in 5 sets of MEFs). Thus, *Keap1-/-* deletions increased viral induction of these transcripts through mechanisms that did not require Nrf2.

To determine if the *Keap1-/-* deletions affected the rate or the efficiency of virus infection or replication, we measured the levels of Sendai virus *M* gene transcripts. The level of *M* transcripts did not increase faster in *Keap1-/- Nrf2-/-* MEFs than in *Nrf2-/-* MEFs (Fig. 1A). The *Keap1-/-* deletions did not increase cytokine gene induction by enhancing Sendai virus infection or replication.

We tested the effects of *Keap1-/-* deletions on *Ifnb1, Tnf* and *Il6* induction by LPS and poly[I:C]. We compared the transcript levels in MEFs with heterozygous *Keap1-/+* and homozygous *Keap1-/-* alleles to distinguish gain and loss of function effects. LPS and poly[I:C] induced 3-6 fold higher *Ifnb1, Tnf*, and *Il6* transcript levels in *Keap1-/- Nrf2-/-* than in *Keap1+/- Nrf2-/-* or in *Keap1+/+ Nrf2-/-* MEFs (Fig. 1B, S1D; range in 2 pairs of MEFs). The *Keap1-/-* deletions increased transcription induction by LPS and poly[I:C] and caused a loss of Keap1 function.

We examined the effects of the *Keap1-/-* deletions on post-induction mechanisms for the regulation of *Ifnb1, Tnf* of *Il6* gene expression. There was no consistent difference in the rates of decline in these transcript levels between *Keap1-/- Nrf2-/-* MEFs and *Nrf2-/-* MEFs beyond 6 h after infection (Fig. 1A, S1A-C). The *Keap1-/-* deletions did not eliminate the post-induction refractory period, and reinfection induced higher levels of *Ifnb1, Tnf* and *Il6* transcripts in *Keap1-/- Nrf2-/-* MEFs than in *Nrf2-/-* MEFs (Fig. 1C).

The *Keap1-/-* deletions had larger effects on *Ifnb1, Tnf* and *Il6* transcript levels in MEFs lacking Nrf2 than in MEFs with intact Nrf2 (Fig. S1A-C). Nrf2 reduced the effects of the *Keap1-/-* deletions through several mechanisms. First, the *Keap1-/-* deletions caused constitutive electrophile response gene transcription in *Keap1-/-* MEFs with intact Nrf2, but not in *Keap1-/- Nrf2-/-* MEFs (Fig. S1A-C; *Nqo1*). Second, the *Keap1-/-* deletions reduced proliferation, and canonical NFκB signaling, in *Keap1-/-* MEFs with intact Nrf2, but not in *Keap1-/- Nrf2-/-* MEFs (Fig. 2C, S2D). Third, Nrf2 counteracted Keap1 binding to the *Ifnb1, Tnf* and *Il6* promoters, and reduced Keap1 effects on the recruitment of other transcription factors and histone modifying enzymes (Fig. 2, 4). We investigated Keap1 functions and the effects of *Keap1-/-* deletions in the presence and in the absence of Nrf2 to distinguish Keap1 effects that are independent of Nrf2 from Keap1 effects that are mediated or modified by Nrf2.

**Fig 2.**
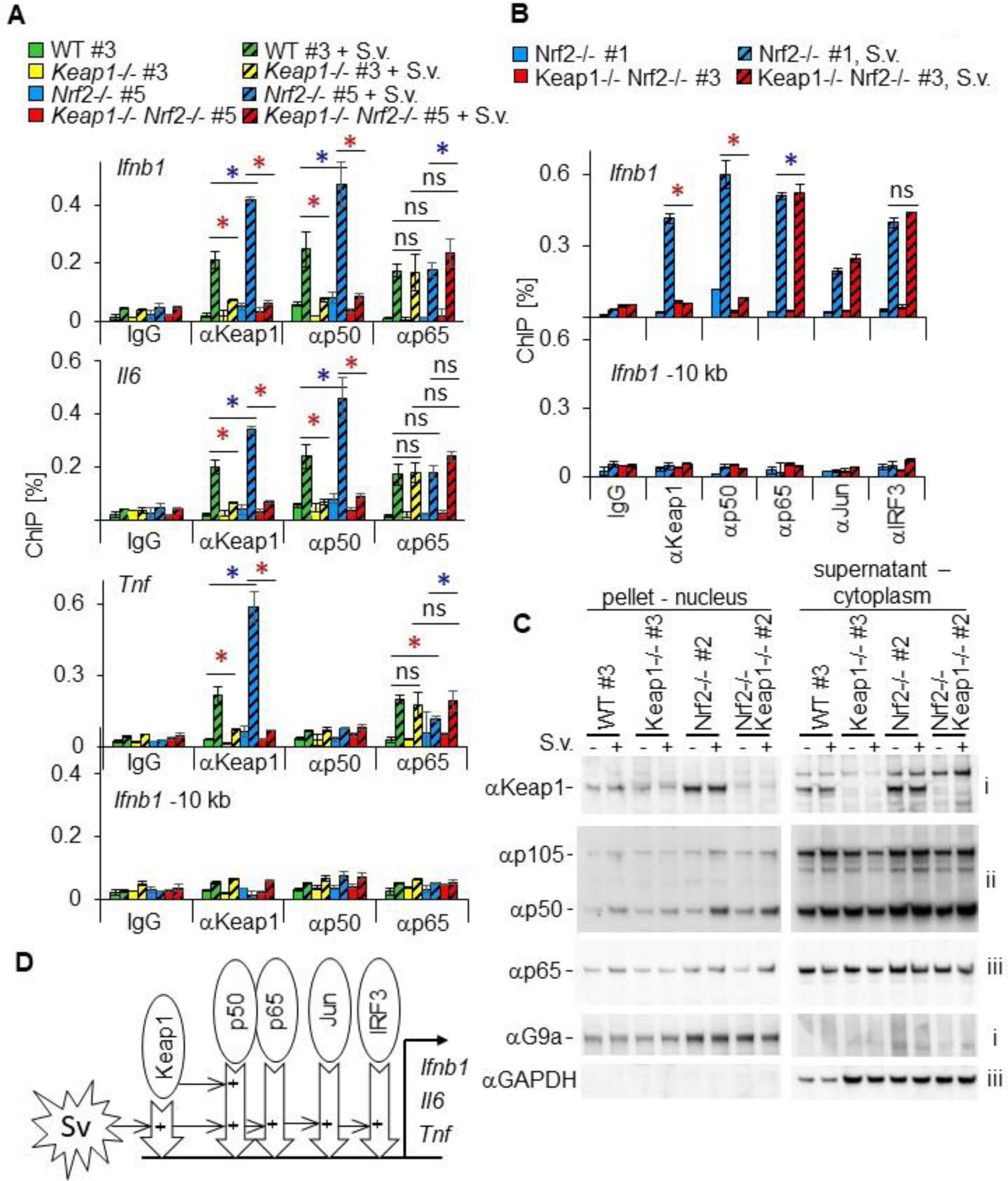
Keap1 binds the *Ifnb1, Tnf*, and *Il6* promoters upon virus infection and is required for NFκB p50 recruitment. (**A**) The levels of Keap1, NFκB p50, and NFκB p65 binding at the *Ifnb1, Tnf*, and *Il6* promoters, and 10 kb upstream of the *Ifnb1* promoter, were measured by ChIP in wild type (WT#3), *Keap1-/- #3, Nrf2-/-#5*, and *Keap1-/- Nrf2-/-#5* MEFs 6 hours after mock (solid bars) and Sendai virus (striped bars) infection. The statistical significance of the differences in the ChIP signals indicated in panels A and B were tested by two-factor ANOVA analyses of data from 2-9 experiments using 2-8 different sets of MEFs (* p<0.001 increase[blue] decrease[red]). The results from analyses of the same samples using αKeap1-C antibodies are shown in Fig. S2B. (**B**) The levels of Keap1, NFκB p50, NFκB p65, cJun and IRF3 binding at the *Ifnb1* promoter and 10 kb upstream of the *Ifnb1* promoter were measured by ChIP in *Nrf2-/-* #1 and in *Keap1-/- Nrf2-/-* #3 MEFs 6h after mock (solid bars) and Sendai virus (striped bars) infection. The results from analyses of binding at the *Tnf, Il6, Ccng* and *Cdkn1a* promoters using the same samples are shown in Fig. S2C. (**C**) The levels and partitioning of Keap1, NFκB p50, NFκB p65, NFκB p105, Gapdh, and G9a in nuclear and cytoplasmic fractions were analyzed following hypotonic lysis of wild type (WT#3), *Keap1-/- #3, Nrf2-/- #2*, and *Keap1-/- Nrf2-/- #2* MEFs after mock (-) or Sendai virus (+) infection as indicated above the lanes. The nuclear pellets (left) and the cytoplasmic supernatants (right) were analyzed in parallel and the antibodies are indicated on the left. The same fractions were analyzed on several membranes as indicated on the right (i, ii, iii). Additional proteins and phosphoproteins in a different set of MEFs are shown in Fig. S2D. (**D**) Effects of Keap1 on transcription factor recruitment to the *Ifnb1, Il6* and *Tnf* promoters. The + symbols inside the hollow arrows indicate the factors on the left that are required for chromatin binding by each protein.

### Keap1 binds to cytokine promoters in response to virus infection

We investigated if Keap1 binds to the *Ifnb1, Tnf* or *Il6* promoters. Since mouse Keap1 binding to specific genes had not been reported, we used antibodies that were raised against three different regions of Keap1 (Fig. S2A, here designated αKeap1, αKeap1-N and αKeap1-C). All three anti-Keap1 antibodies precipitated the *Ifnb1, Tnf*, and *Il6* promoters from virus infected MEFs (Fig. 2A, 2B, S2B, S4A, S4B, 5B and S5B).

We evaluated the specificities of the anti-Keap1 ChIP signals by comparing (1) virus infected *versus* uninfected MEFs, (2) promoter regions *versus* flanking regions, (3) anti-Keap1 *versus* control IgG antibodies, and (4) MEFs with *Keap1-/-* and *Nrf2-/-* deletions. The αKeap1 ChIP-signals at the *Ifnb1, Tnf* and *Il6* promoters were 11 ± 2, 14 ± 5, and 12 ± 2 fold higher, respectively, in virus infected MEFs than in uninfected MEFs (Fig. 2A, 2B, S2C, 5B, S5B, 6A; mean ± sem in 7 MEFs). The αKeap1-C ChIP signals at the same promoters were 24 ± 7, 9 ± 2 and 26 ± 7 fold higher, respectively, in virus infected MEFs than in uninfected MEFs (Fig. S2B, S4A, S5B; mean ± sem in 6 MEFs that partially overlapped the previous set). The αKeap1-N ChIP signal at the *Ifnb1* promoter was 5 ± 1 fold higher in virus infected than in uninfected MEFs (Fig. 4B, S5B; mean ± sem in 2 MEFs). The three anti-Keap1 antibodies produced similar relative ChIP signals at the *Ifnb1, Tnf* and *Il6* promoters when they were used in parallel in the same experiment (Fig. S5B).

Each of the anti-Keap1 antibodies detected low ChIP signals at the *Ccng1* and *Cdkn1a* promoters in uninfected MEFs. Sendai virus infection reduced the αKeap1 ChIP signals at these promoters (Fig. S2C, S5B, S6A; 3 MEFs). Virus infection induced Keap1 binding at the *Ifnb1, Tnf* and *IL6* promoters selectively.

We compared Keap1 binding at the *Ifnb1* promoter and in regions flanking the *Ifnb1* gene. The αKeap1 ChIP signal at the *Ifnb1* promoter was 9 ± 3 fold higher than the ChIP signal 10 kb upstream of the *Ifnb1* promoter upon virus infection (Fig. 2A, 2B, 5B, S5B; mean ± sem in 5 different MEFs). The αKeap1-C ChIP signal at the *Ifnb1* promoter was 7 ± 1 fold higher at the *Ifnb1* promoter than 10 kb upstream of the promoter (Fig S2B, S4A, S5B mean ± sem in 4 MEFs). No Keap1 binding was detected 50 kb from the *Ifnb1* promoter (Fig S4B).

We compared the ChIP signals that were produced by the three anti-Keap1 antibodies with the background signals that were produced by control IgG antibodies. The αKeap1, αKeap1-N, and αKeap1-C antibodies produced 11 ± 3, 10 ± 5, and 22 ± 5 fold higher ChIP signals, respectively, than the control IgG antibodies did at the *Ifnb1* promoter (Fig. 2A, 2B, S2B, 4A, 5B, S5B mean ± sem in 2-6 different MEFs). The αKeap1, αKeap1-N, and αKeap1-C antibodies also precipitated the *Tnf* and *Il6* promoters 3 to 19 fold more efficiently than control IgG antibodies did (Fig. 2A, S2B, 4A, 5B and S5B; range in 2-6 different MEFs). It is exceedingly unlikely that all three antibodies precipitated virus-inducible complexes that were formed by a protein other than Keap1 at the *Ifnb1, Tnf* and *Il6* promoters.

We examined the effects of the *Keap1-/-* deletions on the ChIP signals produced by the three anti-Keap1 antibodies. The αKeap1 ChIP signals were 5 ± 0.7, 7 ± 2 and 4 ± 0.8 fold higher at the *Ifnb1, Tnf* and *Il6* promoters, respectively, in MEFs with intact Keap1 than in MEFs with *Keap1-/-* deletions (Fig. 2A, 2B, S2C, 5B, S5B; mean ± sem in 4 pairs of MEFs). The αKeap1-N ChIP signals at these promoters were 1.3 to 8 fold higher in *Nrf2-/-* MEFs than in *Keap1-/- Nrf2-/-* MEFs (Fig. 4B, S5B; range in 2 pairs of MEFs). By contrast, the *Keap1-/-* deletions did not eliminate the αKeap1-C ChIP signals at the *Ifnb1, Tnf* and *Il6* promoters (Fig. S2B, S4A, S5B). The sequence that encodes the region of Keap1 that was used to raise the αKeap1-C antibodies was not removed by the *Keap1-/-* deletions (Fig. S2A). αKeap1-C antibodies detected ChIP signals only in those chromatin regions where αKeap1 and αKeap1-N antibodies also detected ChIP signals. These and other results suggest that the αKeap1-C antibodies detected chromatin binding by Keap1, and by a fragment of Keap1, a Keap1-related protein, or an antigenically similar protein.

We examined the effects of *Nrf2-/-* deletions on viral induction of Keap1 binding at the *Ifnb1, Tnf* and *IL6* promoters. The αKeap1-C ChIP signals at these promoters were 3 ± 0.4, 4 ± 0.4 and 4 ± 1 fold higher, respectively, in virus infected *Nrf2-/-* MEFs than in wild type MEFs (Fig. S2B, S4A; mean ± sem in 4 pairs of MEFs). The αKeap1 ChIP signals at these promoters were also 2 to 3 fold higher in *Nrf2-/-* MEFs than in wild type MEFs (Fig. 2A; range in 2 pairs of MEFs). The higher αKeap1 and αKeap1-C ChIP signals in *Nrf2-/-* MEFs are likely to reflect higher levels of Keap1 binding since the *Nrf2-/-* deletions did not increase the αNFκB p65 ChIP signals (Fig. 2A, S4A). Consequently, Nrf2 counteracted Keap1 binding at these promoters.

### Keap1 is required for viral induction of NFκB p50 binding

We investigated the effects of *Keap1-/-* and *Nrf2-/-* deletions on chromatin binding by other proteins that were recruited to the *Ifnb1, Il6* and *Tnf* promoters in parallel with Keap1. Sendai virus infection induced NFκB p50 binding at the *Ifnb1* and *Il6* promoters in wild type and in *Nrf2-/-* MEFs (Fig. 2A, 2B). In contrast, virus infection induced little or no NFκB p50 binding at these promoters in *Keap1-/-* or in *Keap1-/- Nrf2-/-* MEFs. The αp50 ChIP signals at the *Ifnb1* and *Il6* promoters were significantly higher in wild type and in *Nrf2-/-* MEFs than in *Keap1-/-* or in *Keap1-/- Nrf2-/-* MEFs following virus infection (Fig. 2A, 2B, S2C, 5C, S5C; 5 sets of MEFs). The *Keap1-/-* deletions did not reduce NFκB p65, IRF3 or cJun binding in the same cells (Fig. 2A, 2B, S2C, 4A, 5C, S5C). Keap1 was also required for NFκB p50 binding at the *Ccng* and *Cdkn1a* promoters in virus infected and in uninfected *Nrf2-/-* MEFs (Fig. S2C, S5C, S6B). Consequently, Keap1 affected the relative amounts of different NFκB subunits that were recruited to promoters in both virus infected and in uninfected MEFs (Fig. 2D).

We compared the effects of *Nrf2-/-* deletions on NFκB p50 binding alone and in combination with *Keap1-/-* deletions. The αp50 ChIP signals at the *Ifnb1* and *Il6* promoters were 2.1 ± 0.3 and 2.1 ± 0.3 fold higher in *Nrf2-/-* MEFs than in wild type MEFs following virus infection (Fig. 2A; mean ± sem in 3 pairs of MEFs). In contrast, the *Nrf2-/-* deletions did not increase NFκB p50 binding in *Keap1-/- Nrf2-/-* MEFs compared with *Keap1-/-* MEFs (Fig. 2A). The epistatic effect of *Keap1-/-* deletions *vis a vie Nrf2-/-* deletions on NFκB p50 recruitment indicates that Nrf2 affected NFκB p50 recruitment indirectly, most likely through interaction with Keap1. The parallel effects of *Nrf2-/-* deletions on Keap1 and on NFκB p50 binding indicate that factors that affect Keap1 binding can regulate NFκB p50 recruitment to cytokine promoters.

### Keap1 partitions between the nucleus and the cytoplasm, and is not required for canonical NFκB signaling

We examined the effects of Sendai virus infection on Keap1 and NFκB expression and partitioning in cell lysates. The αKeap1 antibodies detected a 60 kDa band that partitioned at a 6 : 1 ratio between the cytoplasmic supernatant and the nuclear pellet in cell lysates with intact Keap1 (Fig. 2C). This band was not detected in extracts prepared from *Keap1-/-* or from *Keap1-/- Nrf2-/-* MEFs. Similar Keap1 partitioning was observed in different MEFs (Fig. 2C, S2D). Sendai virus infection did not affect Keap1 expression or its partitioning, suggesting that Keap1 binding to the *Ifnb1, Tnf* and *Il6* promoters was not regulated by changes in Keap1 abundance or localization.

Sendai virus infection increased both NFκB p50 and NFκB p65 partitioning into the nuclear fractions (Fig. 2C). The levels and partitioning of NFκB p50 and NFκB p65 were not significantly different between *Nrf2-/-* and *Keap1-/- Nrf2-/-* MEFs (Fig. 2C, S2D). *Keap1-/-* deletions also had no effect on the ratio between NFκB p50 and its p105 precursor.

We examined the effects of *Keap1-/-* and *Nrf2-/-* deletions on kinases and phosphoproteins that have been implicated in the effects of Keap1 on cytokine transcription (*27, 28*). There was no difference in viral induction of NFκB p65 S536 phosphorylation between *Nrf2-/-* and *Keap1-/- Nrf2-/-* MEFs (Fig. S2D). There was also no difference in IKKβ or IκBα levels, or in IKKα/β or TBK1 phosphorylation between these MEFs. In *Keap1-/-* MEFs, viral induction of IKKα/β phosphorylation, IκBα degradation, and NFκB p50 and NFκB p65 nuclear partitioning were reduced compared with wild-type MEFs. These results differ from the observation that IKKβ is stabilized by Keap1 depletion in other cell types (*27, 28*), and could reflect the slower growth of *Keap1-/-* MEFs. Importantly, there were no differences in the levels of these kinases or phosphoproteins between *Nrf2-/-* and *Keap1-/- Nrf2-/-* MEFs, and there were no differences in the levels of NFκB p65 binding at the *Ifnb1, Tnf* or *Il6* promoters in any of the MEFs (Fig. 2A, 2B, S2C, 5C, S5C). The level of Keap1 was higher in *Nrf2-/-* MEFs than in wild type MEFs, which is consistent with the higher levels of Keap1 binding to the *Ifnb1, Tnf* and *Il6* promoters in *Nrf2-/-* MEFs upon virus infection. Changes in canonical NFκB signaling do not account for the selective effect of the *Keap1-/-* deletions on NFκB p50, and not on NFκB p65, recruitment to the *Ifnb1* and *Il6* promoters.

### Keap1-p50 and Keap1-p65 complex distributions in live cells

To determine if Keap1 formed complexes with NFκB proteins, we visualized Keap1 interactions with NFκB p50 and with NFκB p65 by using bimolecular fluorescence complementation (BiFC) analysis. In this approach, two proteins that are fused to fragments of a fluorescent protein are co-expressed (*46*). If the proteins interact with each other, they can facilitate the association of the fluorescent protein fragments and form a fluorescent complex (Fig. 3E).

**Fig. 3.**
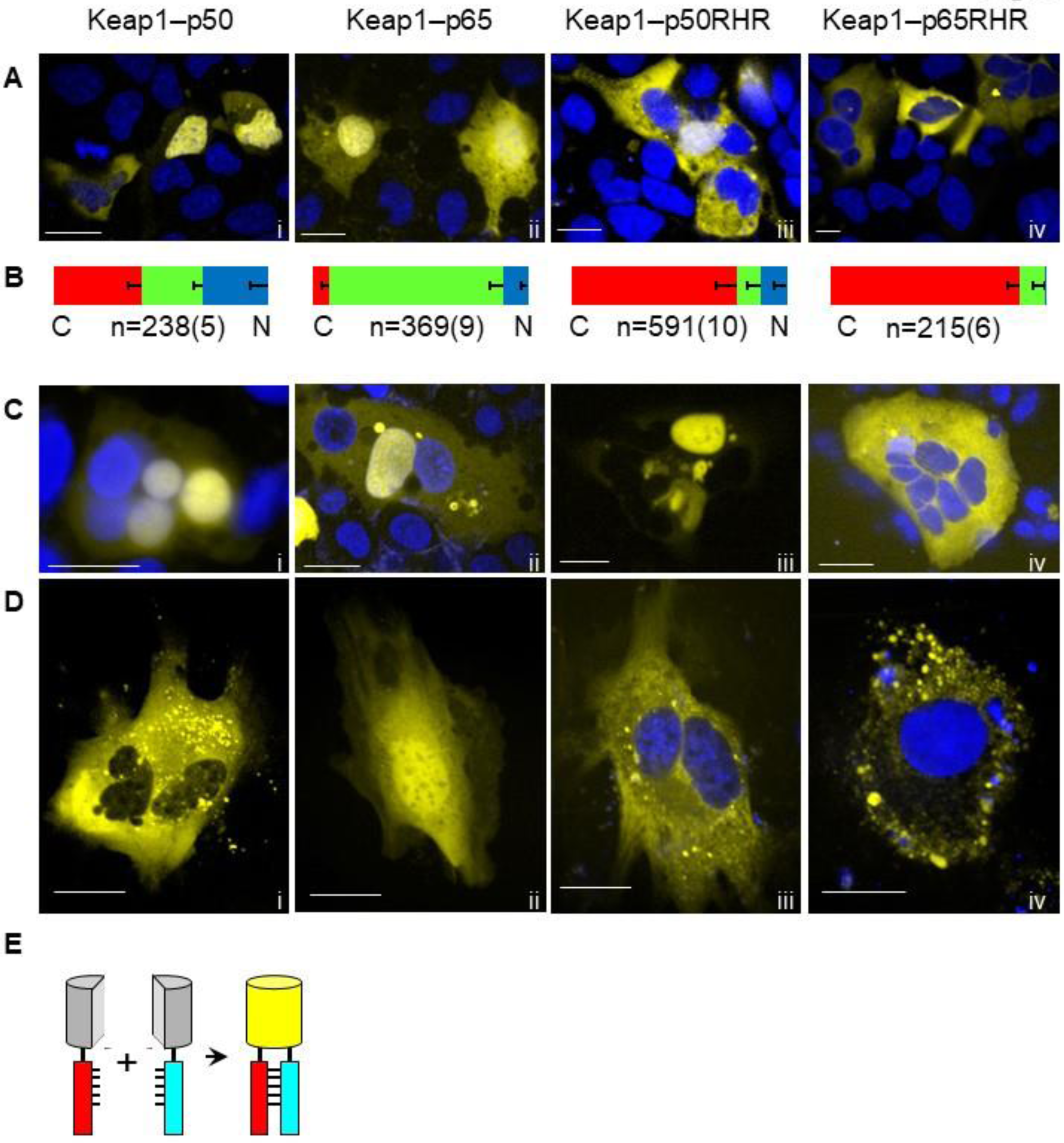
Keap1 forms BiFC complexes with NFκB p50 and NFκB p65 whose subcellular distributions are determined by nuclear factors. (**A**) Each column of images shows cells with fluorescent BiFC complexes formed by the proteins indicated above in HEK293T cells (rows A-D) and in MEFs (row E). The BiFC complex fluorescence is shown in yellow and Hoechst stain is shown in blue. The images were selected to represent the range of different distributions that were observed for each BiFC complex. Scale bars 30 μm. Images of control BiFC complexes that were analyzed in parallel are shown in S3A-C. (**B**) Distributions of the BiFC complexes indicated at the top of each column. The bars show the proportions of cells (mean - sem) in which BiFC complexes were enriched in the cytoplasm (red), the nucleus (blue) and equally in the cytoplasm and the nucleus (green). n: total number of cells (number of experiments in parentheses). (**C**) Images of multinucleate cells with fluorescent BiFC complexes formed by the proteins indicated at the top of each column. The images were selected to represent differences in BiFC complex enrichment among different nuclei within the same multinucleate cell. No Hoechst was added to the cells in image iii. Optical sections of a cell that was imaged at different times are shown in Fig. S3D. (**D**) Fluorescence images of the BiFC complexes indicated at the top of each column in MEFs. The images were selected to show the range of different distributions that were observed for all BiFC complexes that were examined. No Hoechst was added to the cells in images i and ii. (**E**) Diagram of the principle of BiFC analysis (grey - non-fluorescent; yellow - fluorescent).

**Fig. 4.**
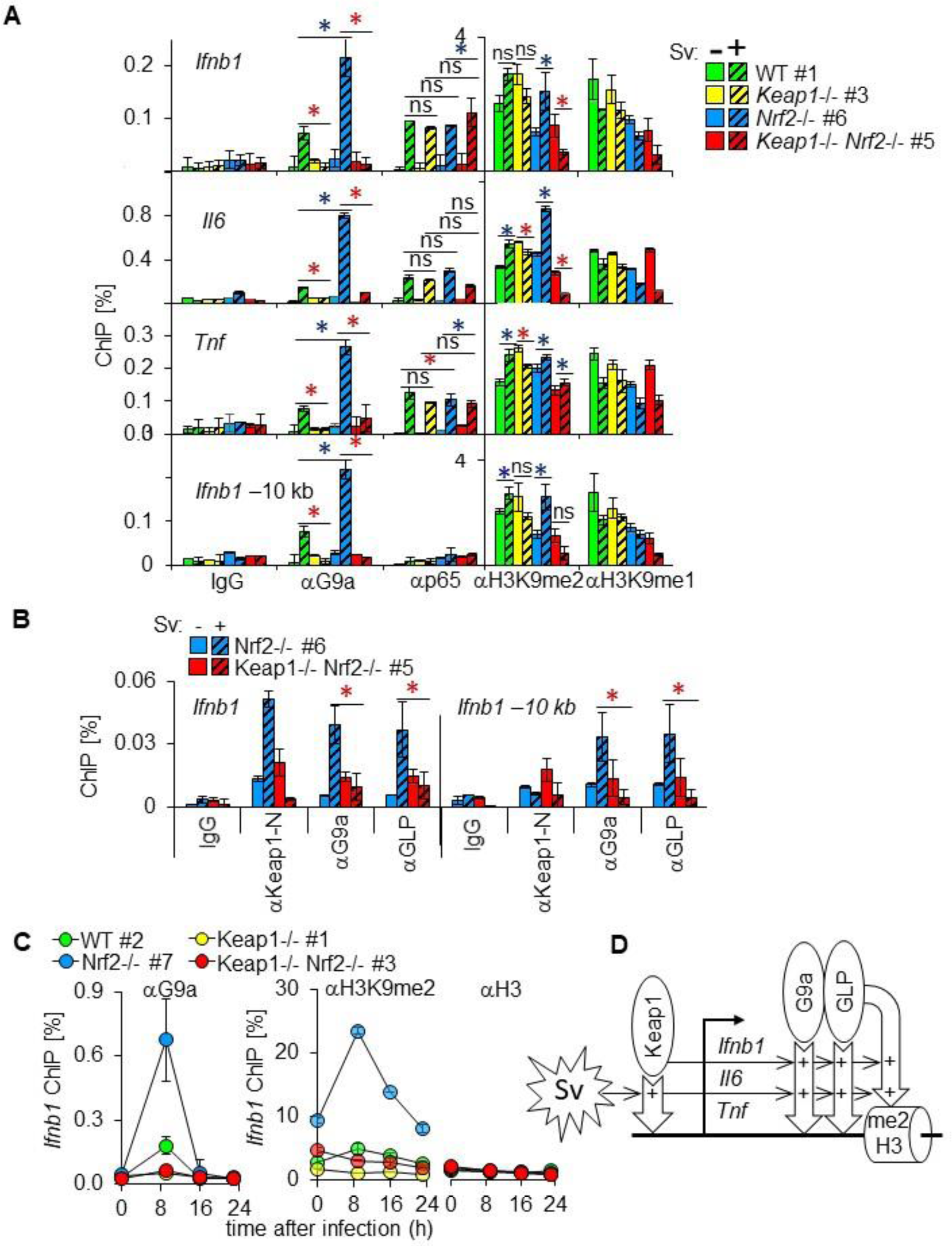
Keap1 is required for G9a-GLP recruitment and H3K9 methylation at the *Ifnb1, Tnf*, and *Il6* genes upon virus infection. (**A**) The levels of G9a and p65 binding, and of H3K9me2 and H3K9me1 were measured at the *Ifnb1, Tnf*, and *Il6* promoters in wild type (WT#1), *Keap1-/- #*3, *Nrf2-/-* #6, and *Keap1-/- Nrf2-/- #*5 MEFs 9 h after mock (solid bars) and Sendai virus (striped bars) infection. A region 10kb upstream of the *Ifnb1* promoter was analyzed in parallel (bottom graphs). The statistical significance of the differences in the ChIP signals indicated in panels A and B were evaluated by two-factor ANOVA analyses of data from 2-9 experiments using 2-8 sets of MEFs (* p<0.001 increase[blue] decrease[red]). Keap1 binding to the same regions, and binding by all proteins to a region 50 kb from the *Ifnb1* promoter was analyzed in parallel using the same samples, and are shown in Fig. S4A and S4B. (**B**) The levels of Keap1, G9a, and GLP binding at the *Ifnb1* promoter and 10 kb upstream of the *Ifnb1* promoter were measured in *Nrf2-/- #*6, and *Keap1-/- Nrf2-/- #*5 MEFs 9 h after mock (solid bars) and Sendai virus (striped bars) infection. (**C**) The levels of G9a binding, H3K9me2 and total H3 at the *Ifnb1* promoter were measured over time following Sendai virus infection of wild type (WT#2), *Keap1-/- #*1, *Nrf2-/- #*7, and *Keap1-/- Nrf2-/- #*4 MEFs. (**D**) Effects of Keap1 on G9a-GLP recruitment and H3K9me2 induction at the *Ifnb1, Il6* and *Tnf* genes by viral infection. The + symbols inside the bent hollow arrow indicate factors that are required for H3K9me2 modification.

We co-expressed Keap1 with NFκB p50, as well as with NFκB p65, fused to complementary fluorescent protein fragments, in HEK293T cells. Both combinations of fusion proteins formed BiFC complexes that had different distributions in different cells (Fig. 3A). Keap1-p50 BiFC complexes were enriched in the nucleus in a third of the cells, in the cytoplasm in a third of the cells, and were equally partitioned between the nucleus and the cytoplasm in a third of the cells (Fig. 3Ai). Keap1-p65 BiFC complexes were equally partitioned between the nucleus and the cytoplasm in a majority of cells, and were enriched in the cytoplasm or in the nucleus in the remaining cells (Fig. 3Aii). In contrast, BiFC complexes formed by Keap1 homodimers and p50-p65 heterodimers had predominantly cytoplasmic and nuclear distributions, respectively, in most cells (Fig. S3A, S3B). The distinct frequencies of Keap1-p50 *versus* Keap1- p65 complex distributions were observed in many independent experiments, suggesting that Keap1 formed separate complexes with NFκB p50 and with NFκB p65 (Fig. 3B).

Keap1 also formed BiFC complexes with fragments encompassing the Rel homology regions (RHR) of NFκB p50 and NFκB p65. These complexes were enriched in the cytoplasm in a majority of cells (Fig. 3Aiii, 3Aiv, 3B). Determinants outside the Rel homology regions of NFκB p50 and NFκB p65 affected the distributions of Keap1-p50 and Keap1-p65 BiFC complexes.

We compared the relative levels of Keap1-p50 as well as Keap1-p65 BiFC complexes among individual nuclei within multinucleate cells. If cytoplasmic factors determine their distributions, then different nuclei are predicted to have equal levels of the BiFC complexes. If nuclear factors determine their distributions, then different nuclei can have different levels of the BiFC complexes. High levels of Keap1-p50 and Keap1-p65 BiFC complexes were present in some nuclei and none were detected in other nuclei within the same multinucleate cells (Fig. 3C). In contrast, Keap1 homodimer and p50-p65 heterodimer BiFC complexes were equally depleted or enriched in all of the nuclei of multinucleate cells (Fig. S3C). The distributions of Keap1-p50 and Keap1-p65 BiFC complexes were generally stable for at least 30 minutes (Fig. S3D). The differences in Keap1-p50 as well as Keap1-p65 BiFC complex levels among nuclei that reside in a common cytoplasm indicate that nuclear factors influence the distributions of these BiFC complexes.

We compared the subcellular distributions of Keap1-p50 and Keap1-p65 BiFC complexes in primary MEFs with those that were observed in HEK293T cells. The distributions of Keap1-p50 and Keap1-p65 BiFC complexes varied among different MEFs, and overlapped with the distributions that were observed in HEK293T cells (Fig. 3D).

### Keap1 is required for G9a-GLP recruitment and for H3K9me2 modification upon virus infection

We investigated if Keap1 affected chromatin modifying enzyme binding or histone modifications at the *Ifnb1, Tnf* or *Il6* genes. Sendai virus infection induced G9a and GLP lysine methyltransferase binding to these genes in wild type MEFs (Fig. 4A, 4B). *Nrf2-/-* deletions enhanced G9a recruitment in MEFs with intact Keap1 (Fig. 4A). Conversely, *Keap1-/-* deletions almost eliminated G9a and GLP recruitment in *Keap1-/-* and in *Keap1-/- Nrf2-/-* MEFs (Fig. 4A, 4B). The αG9a ChIP signals were 7 ± 2, 6 ± 2 and 4 ± 1 fold higher at the *Ifnb1, Tnf* and *Il6* promoters, respectively, in wild type and in *Nrf2-/-* MEFs than in *Keap1-/-* and in *Keap1-/- Nrf2-/-* MEFs (Fig. 4A, 4B, 4C, 5C; mean ± sem in 4 sets of MEFs). Similarly, the αGLP ChIP signals at the *Ifnb1, Tnf* and *Il6* promoters were 3 ± 0.3, 7 ± 4, and 3 ± 0.8 fold higher in *Nrf2-/-* MEFs than in *Keap1-/- Nrf2-/-* MEF (Fig. 4B, 5C; mean ± sem in 2 sets of MEFs). The *Keap1-/-* deletions did not reduce NFκB p65 binding, and the *Nrf2-/-* deletions did not increase NFκB p65 binding at these promoters in the same cells (Fig. 2A, 4A, 5C).

Keap1 was also required for G9a and GLP recruitment to regions 10 kb and 50 kb from the *Ifnb1* promoter (Fig. 4A, S4B). The *Keap1-/-* and *Nrf2-/-* deletions did not affect G9a expression or its nuclear localization (Fig. 2C). The concerted effects of *Keap1-/-* as well as *Nrf2-/-* deletions on Keap1, NFκB p50 and G9a recruitment to several genes suggest that Keap1 binding reconfigured both nucleoprotein complexes and chromatin organization at these genes (Fig. 4D).

We investigated the effects of Sendai virus infection, and of *Keap1-/-* and *Nrf2-/-* deletions, on histone H3 mono- and di-methylation (H3K9me1 and H3K9me2) at the *Ifnb1, Tnf* and *Il6* promoters. Sendai virus infection tended to increase H3K9me2 at these promoters in wild type and in *Nrf2-/-* MEFs (Fig. 4A). Conversely, virus infection generally reduced H3K9me2 at the *Ifnb1* and *IL6* promoters in *Keap1-/-* and in *Keap1-/- Nrf2-/-* MEFs, and reduced or produced a smaller increase in H3K9me2 at the *Tnf* promoter (Fig. 4A, 4C, 5C, S5C). Virus infection induced similar changes in H3K9me2 10 kb upstream of the *Ifnb1* promoter and at the *Ccng1* and *Cdkn1a* promoters (Fig. 4A, S6A). These effects of *Keap1-/-* and *Nrf2-/-* deletions on the changes in H3K9me2 levels following virus infection are consistent with the essential role of Keap1 in G9a and GLP recruitment to these genes.

We measured the levels of G9a binding and of H3K9me2 at different times after infection to determine if the *Nrf2-/-* deletions affected the persistence of G9a binding or H3K9me2 modification at the *Ifnb1* promoter. Both G9a binding and H3K9me2 modification were transient and returned to baseline levels within 16 and 23 h, respectively (Fig. 4C). The *Nrf2-/-* deletions increased the magnitudes of the responses, but did not alter their timing.

### G9a-GLP inhibitors reverse transcription repression by Keap1

We investigated the effects of the G9a-GLP inhibitor BIX01294 on viral induction of *Ifnb1, Il6* and *Tnf* transcription to analyze the role of lysine methyltransferase activity in their repression by Keap1. The addition of BIX01294 one hour before virus infection increased the peak *Ifnb1, IL6* and *Tnf* transcript levels significantly more in *Nrf2-/-* MEFs than in *Keap1-/- Nrf2-/-* MEFs (Fig. 5A, S5A; 4 sets of MEFs). BIX01294 did not increase the levels of Sendai virus *M* gene transcripts. The rapid and selective enhancement of virally induced transcription by BIX01294 in *Nrf2-/-* MEFs, but not in *Keap1-/- Nrf2-/-* MEFs is consistent with the hypothesis that Keap1 represses transcription by recruiting G9a-GLP.

**Fig. 5.**
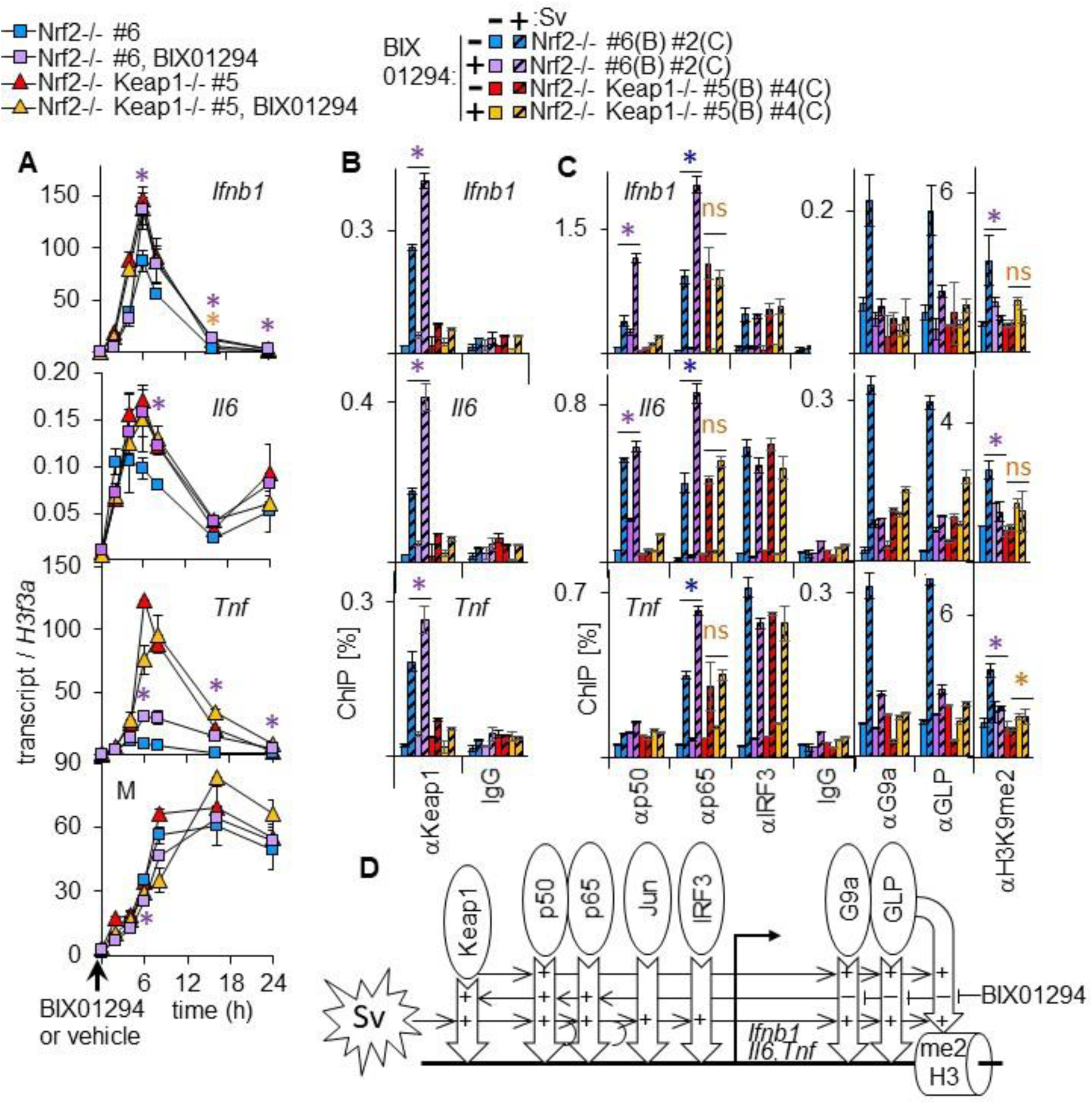
BIX01294 enhances viral induction of transcription, and Keap1 and NFκB recruitment, selectively in MEFs with intact Keap1. (**A**) The levels of *Ifnb1, Il6, Tnf*, and *M* gene transcripts were measured in *Nrf2-/- #*6, and in *Keap1-/- Nrf2-/- #*5 MEFs at the indicated times after Sendai virus infection following 20 μM BIX01294 or vehicle addition 1 h before infection (arrow on time axis). The statistical significance of the effects of BIX01294 on transcription were tested by two-factor ANOVA analyses of data from 2 experiments with different pairs of MEFs (* p<0.0001 *Nrf2-/-* [purple] *Keap1-/- Nrf2-/-* [orange]). The effects of BIX01294 on transcription in different MEFs are shown in Fig. S5A. (**B**) The levels of Keap1 binding at the *Ifnb1, Il6*, and *Tnf* promoters were measured in *Nrf2-/-* #6, and *Keap1-/- Nrf2-/- #*5 MEFs 6 h after mock (solid bars) and Sendai virus (striped bars) infection following 20 μM BIX01294 or vehicle addition one hour before infection. The statistical significance of the effects of BIX01294 on the ChIP signals indicated in panels B and C were tested by two-factor ANOVA analyses of data from 3 experiments using different pairs of MEFs (* p<0.0005 *Nrf2-/-* [purple] *Keap1-/- Nrf2-/-* [orange]). Analyses of NFκB p50 and NFκB p65 binding and H3K9me2 in the same samples are shown in Fig. S5C. (**C**) The levels of NFκB p50, NFκB p65, IRF3, G9a, and GLP binding, and H3K9me2 at the *Ifnb1, Il6 and Tnf* promoters were measured in *Nrf2-/-* #2, and *Keap1-/- Nrf2-/- #*4 MEFs 6 h after mock (solid bars) and Sendai virus (striped bars) infection following 20 μM BIX01294 or vehicle addition one hour before virus infection. Keap1 binding was analyzed in parallel using the αKeap1, αKeap1-N, and αKeap1-C antibodies, and is shown in Fig. S5B. (**D**) Effects of Keap1 and of BIX01294 on transcription factor and lysine methyltransferase recruitment to the *Ifnb1, Il6* and *Tnf* promoters. The + symbols inside the hollow arrows indicate factors that are required or enhance, and the – symbols indicate factors that inhibit, chromatin binding or modification.

### G9a-GLP inhibitors enhance viral induction of Keap1 and NFκB binding

We investigated the effects of BIX01294 on Keap1 recruitment and of *Keap1-/-* deletions on BIX01294 effects on the recruitment of other proteins. BIX01294 increased all anti-Keap1 ChIP signals when it was added to *Nrf2-/-* MEFs one hour before Sendai virus infection. The αKeap1 ChIP signals at the *Ifnb1, Tnf* and *Il6* promoters were significantly higher in virus infected *Nrf2-/-* MEFs that were cultured with BIX01294 than in equivalent MEFs that were cultured with vehicle (Fig. 5B, S5B; 3 MEFs). The αKeap1-N and αKeap1-C ChIP signals at the *Ifnb1, Tnf* and *IL6* promoters were also higher in virus infected *Nrf2-/-* MEFs that were cultured with BIX01294 than in equivalent MEFs that were cultured with vehicle (Fig. S5B).

BIX01294 increased the αp65 and αp50 ChIP signals at the *Ifnb1, Il6*, and *Tnf* promoters in *Nrf2-/-* MEFs. The αp65 ChIP signals at the *Ifnb1, Il6* and *Tnf* promoters were significantly higher in virus infected *Nrf2-/-* MEFs that were cultured with BIX01294 than in equivalent MEFs that were cultured with vehicle. The αp50 ChIP signal at the *Ifnb1*, promoter was significantly higher in the same MEFs (Fig. 5C, S5C; 3 pairs of MEFs for both αp65 and αp50). In contrast, BIX01294 did not increase NFκB p65 or p50 binding in *Keap1-/- Nrf2-/-* MEFs (Fig. 5C, S5C). BIX01294 also did not alter the αIRF3 or the αH3 ChIP signals at any of the promoters in any of the MEFs that were tested (Fig. 5C, S5C). BIX01294 increased NFκB p50 and NFκB p65 binding selectively in *Nrf2-/-* MEFs with intact Keap1, and not in *Keap1-/- Nrf2-/-* MEFs (Fig. 5D).

BIX01294 inhibited G9a and GLP recruitment to all of the promoters and flanking regions that were examined (Fig. 5C). BIX01294 also inhibited the increase in H3K9me2 that was induced by virus infection in *Nrf2-/-* MEFs that were cultured with vehicle (Fig. 5C, S5C). The opposite effects of BIX01294 on Keap1, NFKB p50 and NFKB p65 *versus* G9a and GLP recruitment in the same cells corroborate the selectivity of BIX01294 effects. The elimination of G9a and GLP recruitment to the *Ifnb1, Il6*, and *Tnf* genes both by *Keap1-/-* deletions and by culture with BIX01294 correlated with greater viral induction of their transcription under both circumstances.

### Several lysine methyltransferase inhibitors enhance viral induction of Keap1 and NFκB p50 binding and of transcription

Inhibition of lysine methyltransferase activity by structurally distinct G9a-GLP inhibitors (BIX01294, MS012) increased Keap1 and NFκB p50 recruitment to the *Ifnb1* and *Il6* promoters in *Nrf2-/-* MEFs (Fig. 6A). Each compound increased NFκB p50 binding in *Nrf2-/-* MEFs, but did not induce NFκB p50 binding *Keap1-/- Nrf2-/-* MEFs. Likewise, each compound inhibited viral induction of H3K9me2 at the *Ifnb1* and *Il6* promoters in *Nrf2-/-* MEFs, but did not reduce H3K9me2 in *Keap1-/- Nrf2-/-* MEFs. The compounds had little or no effect on H3K27me3 at the *Ifnb1* and *Il6* promoters in either of the MEFs (Fig. S6A). The GSK343 inhibitor of Ezh2 reduced H3K27me3 at the *Ifnb1* and *Il6* promoters, and had smaller effects on H3K9me2 in both MEFs (Fig. S6A). It is likely that the higher levels of Keap1 and NFκB p50 binding in *Nrf2-/-* MEFs that were cultured with BIX01294 and MS012 were due to the inhibition of G9a-GLP lysine methyltransferase activity.

**Fig. 6.**
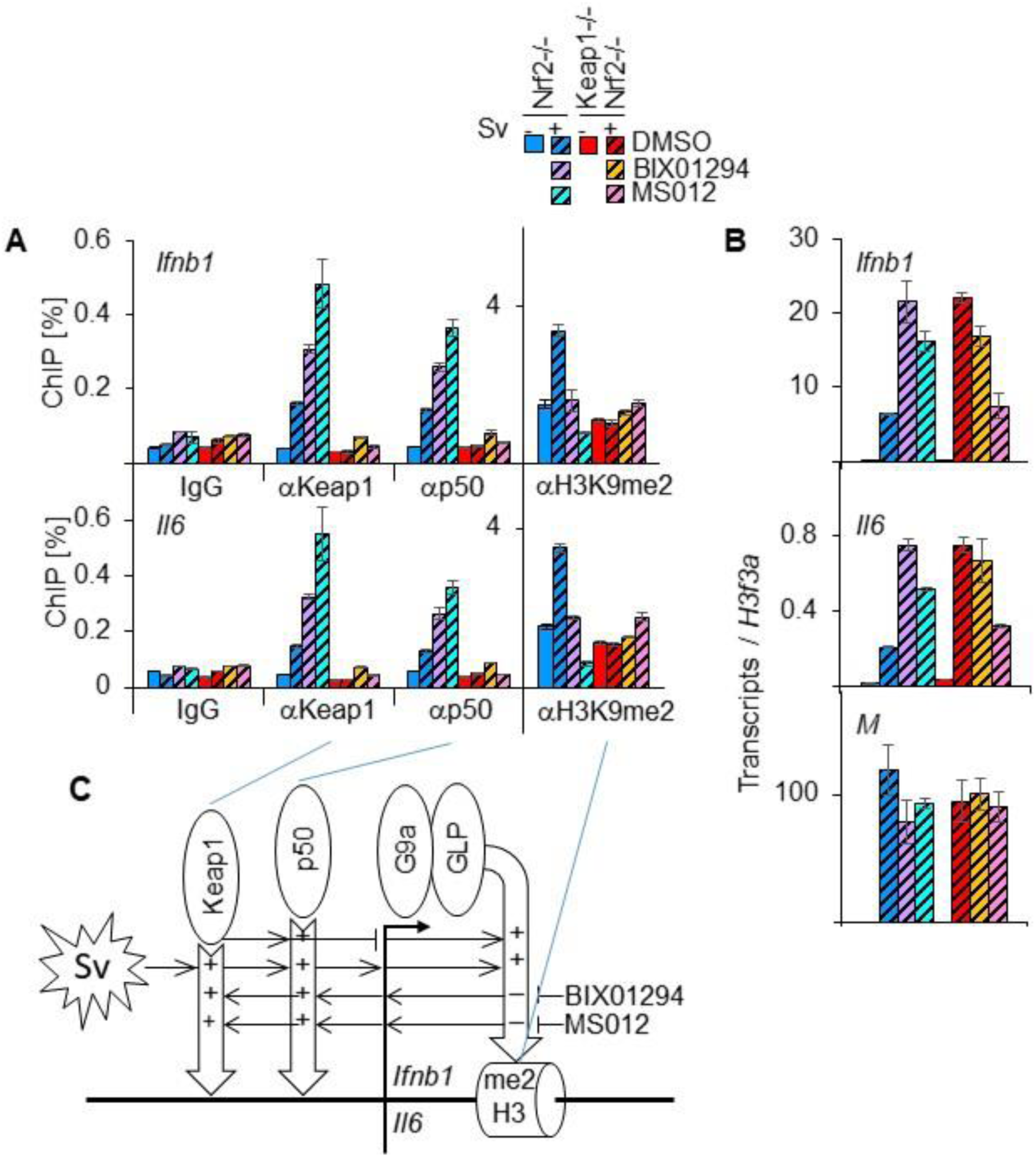
Different lysine methyltransferase inhibitors enhance viral induction of Keap1 and NFκB p50 binding, and *Ifnb1* and *Il6* transcription, selectively in MEFs with intact Keap1. (**A**) The levels of Keap1, NFκB p50, and H3K9me2 binding at the *Ifnb1* and *Il6* promoters were measured in *Nrf2-/-* #6 and *Keap1-/- Nrf2-/- #*5 MEFs 6 hours after mock (solid bars) and Sendai virus (striped bars) infection following 1 h culture with 20 μM BIX01294, or 48 h culture with 1 μM MS012. (**B**) The levels of *Ifnb1, Il6*, and *M* transcripts were measured in *Nrf2-/- #*6 and in *Keap1-/- Nrf2-/- #*5 MEFs that were cultured in parallel with the MEFs that were analyzed in panel A. Each graph in this Fig. shows the means and the standard errors of duplicate samples from one experiment. (**C**) Effects of different lysine methyltransferase inhibitors on Keap1 and NFκB p50 binding, on H3K9me2, and on *Ifnb1* and *Tnf* transcription.

Both BIX01294 and MS012 increased viral induction of *Ifnb1* and *Il6* transcription in *Nrf2-/-* MEFs (Fig. 6B). In contrast, they reduced, or had little effect, on *Ifnb1* and *Il6* transcription in *Keap1-/- Nrf2-/-* MEFs. Virus infection induced higher levels of *Ifnb1* and *Il6* transcription in *Keap1-/- Nrf2-/-* MEFs than in *Nrf2-/-* MEFs, consistent with the capacity of Keap1 to moderate viral induction of transcription in the absence of lysine methyltransferase inhibitors. GSK343 had a small effect on *Ifnb1* and *Il6* transcription in *Nrf2-/-* MEFs, but increased their transcription in *Keap1-/- Nrf2-/-* MEFs (Fig. S6B). The effects of these compounds on viral induction of transcription were consistent with the hypothesis that the ability of Keap1 to moderate *Ifnb1* and *Il6* induction by virus infection requires G9a-GLP lysine methyltransferase activity (Fig. 6C).

## Discussion

Keap1 binding to the *Ifnb1, Tnf* and *Il6* promoters moderates their induction by Sendai virus infection. Three sets of observations corroborate Keap1 regulation of these genes: (1) Keap1 is essential for NFκB p50 and G9a-GLP recruitment to these genes, (2) Keap1 affects NFκB p50, but not p65 recruitment, and (3) lysine methyltransferase inhibitors enhance viral induction of transcription, and Keap1 and NFκB recruitment.

Keap1 is likely to regulate many immunomodulatory genes because: (a) The *Ifnb1, Tnf* and *Il6* promoters are activated by different mechanisms (*44, 45*), suggesting that Keap1 can moderate transcription activation by several mechanisms; (b) NFκB p50 and G9a-GLP can bind and regulate many immune response genes (*4, 14*), suggesting that Keap1 can control NFκB dimer composition and chromatin modification at these genes; and (c) Keap1 depletion in mice and in cultured cells alters immune regulation and cytokine induction (*27-29, 35, 36*), suggesting that Keap1 modulates immune functions in many cell types.

Keap1 does not contain sequence similarity to known DNA or histone binding domains. Keap1 is likely to be recruited to promoters through interactions with DNA or chromatin binding proteins. It is notable that Keap1 binding is more selective than binding by the established transcription regulators at the genes that we tested. It is possible that a combination of positive and negative effects by several of the proteins that are co-recruited with Keap1 determines the selectivity of Keap1 recruitment. It is also possible that additional determinants contribute to the specificity of Keap1 recruitment by virus infection.

Keap1 forms complexes with NFκB p50 and with NFκB p65 in live cells. The differences in Keap1-p50 and Keap1-p65 BiFC complex characteristics suggest that Keap1 forms separate complexes with NFκB p50 and NFκB p65. The distinct characteristics of Keap1-p50 and Keap1-p65 BiFC complexes correlate with the distinct effects of *Keap1-/-* deletions on NFκB p50 *versus* NFκB p65 recruitment to promoters upon virus infection. These observations provide insight into the molecular mechanisms that mediate the selective recruitment of NFκB p50 to genes whose expression is regulated by immunomodulatory signals (*6, 14*).

Virus infection induces Keap1 binding to the *Ifnb1, Tnf* and *Il6* promoters and G9a-GLP binding throughout the *Ifnb1* gene. Keap1 is required for G9a-GLP recruitment and for increased H3K9me2 both at the promoter and in flanking regions. Long-range effects of Keap1 binding at the promoters could be mediated by mechanisms that involve contacts between distant chromatin regions or spreading along the chromatin fiber. Since a basal level of H3K9me2 is present in the absence of virus infection, it is likely that Keap1 facilitates G9a-GLP occupancy in flanking regions by mechanisms in addition to H3K9me2 spreading.

Keap1 has parallel effects on NFκB p50 and on G9a-GLP binding at the *Ifnb1* and *Il6* promoters. This is consistent with the previous observation that NFκB p50 and GLP coprecipitate from HEK293T cell extracts (*3*). Whereas NFκB p50 and G9a-GLP are co-recruited to some genes by Keap1, they can also be recruited separately, and lysine methyltransferase inhibitors have opposite effects on Keap1 recruitment of NFκB p50 *versus* G9a-GLP.

The rapid and selective derepression of *Ifnb1, Tnf* and *Il6* transcription by BIX01294 in MEFs with intact Keap1, but not in MEFs lacking Keap1, indicates that Keap1 represses transcription through mechanisms that require lysine methylation. The selective enhancement of Keap1 and NFκB recruitment and viral induction of transcription by BIX01294 only in MEFs with intact Keap1, corroborates the interrelationship between G9a-GLP activity on the one hand and Keap1 and NFκB recruitment on the other. The reciprocal effects of Keap1 and of G9a-GLP on chromatin binding by each other creates a feedback circuit that modulates their occupancy in response to signals that alter either Keap1 recruitment or G9a-GLP activity. The rapid effects of BIX01294 on promoter occupancy and activity corroborate the effects of *Keap1-/-* and *Nrf2-/-* deletions on transcription factor recruitment and viral induction of transcription. Whereas *Keap1-/-* and *Nrf2-/-* deletions affect mainly NFκB p50 recruitment, BIX01294 enhances both NFκB p50 and NFκB p65 recruitment, indicating that BIX01294 is likely to influence their recruitment through mechanisms in addition to increased Keap1 binding.

The effects of *Keap1-/-* deletions on many immune functions have been interpreted to be due to Nrf2 activation (*25, 34-37*). In contrast to this interpretations, we found that *Nrf2-/-* deletions influence NFκB p50 and G9a-GLP recruitment by modulating Keap1 binding at the *Ifnb1, Tnf* and *Il6* promoters. Indirect effects of Keap1 on cytokine gene regulation, including the destabilization of IKKβ, can also contribute (*27, 28*). Physiological states as well as pharmacological agents that shift the balance among these effects are likely to affect immune responsiveness. Keap1 regulation of cytokine induction is a tractable target for the development of new compounds that can modulate innate immune responses.

## Materials and Methods

### Mice

Mice that carried germline deletions in the genes that encode Keap1 and Nrf2 were generated by the Masayuki Yamamoto laboratory, and were obtained from the Thomas Kenzler laboratory (*47, 48*). These mice were crossed to generate *Keap1-/+* and *Nrf2-/+* double heterozygous stocks. These stocks were crossed to produce embryos with different combinations of Keap1 and Nrf2 alleles. *Nrf2-/-* embryos were generated by crossing homozygous *Nrf2-/-* mice. *Keap1+/+ Nrf2+/+* embryos were generated by crossing C57Bl/6 mice. The mice were housed under specific pathogen-free conditions and experimental protocols were approved by the University of Michigan Animal Care and Use Committee.

### MEF derivation

Mouse embryo fibroblasts (MEFs) that carried deletions in the genes encoding Keap1 and/or Nrf2 (designated *Keap1-/-* and *Nrf2-/-*) were derived from embryos isolated at about 13 days after coitus. MEFs of each genotype were obtained from several different litters. Each MEF preparation is identified by a number following # in the figures and legends.

### Transcript Analyses

MEFs of the indicated genotypes were cultured in parallel. In some experiments, compounds or vehicle were added at the indicated times before virus infection. Sendai virus (Charles River Laboratories, Wilmington, MA) was added at 200 hemagglutinin units/ml. RNA was isolated and the indicated transcript levels were measured by RT-qPCR.

### ChIP Analyses

MEFs were cultured as described above, crosslinked, and the chromatin was sheared by sonication. Transcription factors and histones were precipitated using the antibodies indicated, and the levels of co-precipitated promoter and flanking regions were measured by ChIP-qPCR.

### Partitioning into nuclear and cytoplasmic fractions

MEFs were cultured as described above, lysed, and the nuclei were separated from the cytoplasm by centrifugation. The relative levels of the proteins indicated were measured by immunoblotting.

### BiFC analyses of protein interactions in living cells

Plasmids that encoded proteins in which Keap1 and NFκB p50 or NFκB p65 were fused to complementary YFP fragments (YN and YC), were transfected into HEK293T cells and MEFs. The subcellular distributions of BiFC complexes formed by the fusion proteins were imaged by fluorescence microscopy in living cells that were stained using Hoechst.

### Materials

A complete list of the materials that were used is provided in supplementary materials. The specificities of the antibodies were evaluated by immunoblotting and by parallel experiments using *Keap1-/-* MEFs.

### Statistical analyses

Two-factor ANOVA with replication was used to analyze data from independent experiments that were performed with MEFs from different embryos.

## Acknowledgments

We thank Masayuki Yamamoto for developing, and Thomas Kenzler for providing, mouse strains with the *Keap1* and the *Nrf2* deletion alleles. We thank Huai Deng for isolating some of the MEFs and for sharing his results from experiments to identify genes that were bound and regulated by Keap1 in MEFs.

## Author contributions

TKK conceived the study. VEB and TKK designed the experiments. VEB performed the RT-qPCR and ChIP experiments. TKK performed the BiFC experiments. TKK supervised the experiments and wrote the paper.

## List of Supplementary Materials

The following information and data are available as supplementary materials in a separate file: Supplementary Materials and Methods

Fig. S1 *Keap1-/-* deletions increase *Ifnb1, Tnf*, and *Il6* induction in multiple independently derived MEFs lacking Nrf2, and increase *Nqo1* transcription only in MEFs with intact Nrf2.

Fig. S2 Virus infection induces Keap1 binding at the *Ifnb1, Tnf*, and *Il6* promoters, but not at the *Ccng* or *Cdkn1* promoters. Keap1 is required for NFκB p50, but not for NFκB p65 recruitment, or for canonical NFκB signaling.

Fig. S3 Keap1-Keap1, p50-p65, p50-p65RHR, Keap1-p65 and Keap1-p50RHR BiFC complexes differ in subcellular distributions, and in differential enrichment among individual nuclei within multinucleate cells.

Fig. S4 Keap1 is required for G9a recruitment and for H3K9 methylation throughout the *Ifnb1* locus upon virus infection.

Fig. S5 BIX01294 enhances virus induction of transcription and Keap1 and NFκB binding at cytokine promoters selectively in MEFs with intact Keap1.

Fig. S6 G9a-GLP lysine methyltransferase inhibitors selectively enhance Keap1 and NFκB p50 binding in *Nrf2-/-* MEFs.

References (49-52)

