## Supplementary Figures, Materials and Methods for "Keap1 binds cytokine promoters upon virus infection and moderates their induction by recruiting NFκB p50 and G9a-GLP"

**Supplementary Materials Table of Contents**


| | | |
|---|---|---|
| Supplementary Materials and Methods | | 2 |
| Fig. S1 | *Keap1-/-* deletions increase *Ifnb1*, *Tnf*, and *Il6* induction in multiple independently derived MEFs lacking Nrf2, and increase *Nqo1* transcription only in MEFs with intact Nrf2. | 9 |
| Fig. S2 | Virus infection induces Keap1 binding at the *Ifnb1*, *Tnf*, and *Il6* promoters, but not at the *Ccng* or *Cdkn1* promoters. Keap1 is required for NFκB p50, but not for NFκB p65 recruitment, or for canonical NFκB signaling. | 11 |
| Fig. S3 | Keap1-Keap1, p50-p65, p50-p65RHR, Keap1-p65 and Keap1-p50RHR BiFC complexes differ in subcellular distributions, and in differential enrichment among individual nuclei within multinucleate cells. | 14 |
| Fig. S4 | Keap1 is required for G9a recruitment and for H3K9 methylation throughout the *Ifnb1* locus upon virus infection. | 16 |
| Fig. S5 | BIX01294 enhances virus induction of transcription and Keap1 and NFκB binding at cytokine promoters selectively in MEFs with intact Keap1. | 18 |
| Fig. S6 | G9a-GLP lysine methyltransferase inhibitors enhance Keap1 and NFκB p50 binding in *Nrf2-/-* MEFs selectively | 20 |

**Supplementary Materials and Methods**

**Mice.** Mice that carried germline deletions in the genes encoding Keap1 and/or Nrf2 were generated by the Masayuki Yamamoto laboratory, and were obtained with permission from the Thomas Kenzler laboratory (*42*, *43*). These mice were crossed to generate *Keap1-/+* and *Nrf2-/+* double heterozygous stocks. These stocks were crossed to produce embryos with different combinations of Keap1 and Nrf2 alleles. *Nrf2-/-* embryos were generated by crossing homozygous *Nrf2-/-* mice. *Keap1+/+ Nrf2+/+* embryos were generated by crossing C57Bl/6 mice. The mice were housed under specific pathogen-free conditions and experimental protocols were approved by the University of Michigan Animal Care and Use Committee.

**MEF derivation and culture conditions.** Mouse embryo fibroblasts (MEFs) that carried deletions in the genes encoding Keap1 and/or Nrf2 (here designated *Keap1-/-* and *Nrf2-/-*) were derived from embryos isolated at about 13 days after coitus. MEFs of each genotype were obtained from several different litters. The genotypes of the embryos were determined by PCR analyses as described (*47*, *48*). Each MEF preparation is identified by a number following # in the figures and legends. MEFs from different litters were compared in each experiment to eliminate the possibility that the differences between the MEFs were unique to a particular litter. The MEFs were cultured in DMEM (GIBCO, cat. no. 11995) supplemented with 10% FBS and 0.1 mg/ml penicillin-streptomycin. MEFs stocks were obtained after cell expansion to passage 2 or 3. Passage 3, 4 and 5 MEFs were used for all experiments.

**MEF culture and Sendai virus infection.** MEFs of the indicated genotypes were plated at a density of 200,000/ml. MEFs for transcript analysis and cell extract fractionation were cultured in 6-well plates. MEFs for ChIP analyses were cultured in 15 cm plates according to standard procedures (*49*). In experiments that included culture with different compounds, the compounds were added to the cultures at the concentrations and times indicated before virus infection. The MEFs were infected with Sendai virus by the addition of 200 hemagglutinin units/ml of Sendai virus (Charles River Laboratories, Wilmington, MA). MEFs of the indicated genotypes were cultured in parallel and were harvested at the indicated times after virus and/or inhibitor addition. For time course analyses of Sendai virus infection, aliquots of the same Sendai virus stock were thawed and added in an identical manner at different times and the cells were harvested and analyzed in parallel. LPS (Sigma L4391) was dissolved in PBS (Gibco 100100) to a stock concentration of 20 μg/ml and frozen aliquots. Aliquots were thawed immediately before addition to cultured cells at a final concentration of 200 ng/ml. Poly(I:C) (Tocris 4287) was dissolved in RNase-free water included with Roche First Strand cDNA Synthesis Kit (04379012001) at a stock concentration of 400 μg/ml and frozen. Aliquots were thawed immediately before addition to cultured cells at a final concentration of 4 μg/ml.

To test the effects of tBHQ or BIX01294 on viral induction of cytokine transcription or protein binding, aliquots of the same stock of either tBHQ or BIX01294 were added at a final concentrations of 50 μM (tBHQ) or 20 μM (BIX01294) either 24 h (tBHQ) or 1 h (BIX01294) before Sendai virus addition. Vehicle (DMSO or ddH2O) was added to control MEFs.

**Analysis of transcript levels by RT-qPCR.** The MEFs (200,000 cells per sample) were washed twice with pre-warmed 1X PBS (Gibco cat. no 10010023), released with 0.05% trypsin-EDTA (Gibco cat no. 25300054), and suspended in culture medium. The cells were collected by 5 min centrifugation at 2000 x g, the supernatant was removed, and 350 µl of RLT buffer (QIAgen RNeasy Kit) supplemented with 1% (v/v) β-mercaptoethanol was added. mRNA was isolated using the RNeasy Mini Kit (Qiagen). Equal amounts of mRNA from all samples were reverse transcribed using the Transcriptor First Strand cDNA synthesis kit under conditions recommended by the manufacturer (Roche). The relative amounts of cDNAs corresponding to the indicated transcripts were measured using qPCR assays. Each qPCR reaction contained 4% (vol/vol) cDNA, 1 µM of each primer, and 1X Roche SYBR Green Master Mix. The relative transcript levels were calculated by assuming that they were inversely proportional to $2^{Ct}$, where $C_t$ equals the number of cycles required to reach threshold fluorescence. The amounts of transcripts in each sample were normalized by the amount of *H3f3a* transcripts in the same sample. The average normalized transcript level for duplicate reactions was plotted with upper and lower error bars corresponding to 2 times the standard deviation. The oligonucleotide primers for RT-qPCR were obtained from IDT, were validated by melt-curve analyses, and had the following sequences:

| **RT-qPCR** | **Forward primer sequence** | **Reverse primer sequence** |
|---|---|---|
| *Ifnb1* | cac agc cct ctc cat caa cta | Cat ttc cga atg ttc gtc ct |
| *Tnf* | Ttg tct taa taa cgc tga ttt ggt | Ggg agc aga ggt tca gtg at |
| *Il6* | Tgc ctt cat tta tcc ctt gaa | Tta cta cat tca gcc aaa aag cac |
| M | Tgg tgc tcc act cct acc at | Gtg cga cct tgt ttg cat ta |
| *Ccng1* | Cac acc aat cag cga cgt a | Ggg aac aag ctg gag acc t |
| *Cdkn1a* | Tcc aca gcg ata tcc aga ca | gga cat cac cag gat tgg ac |
| *H3f3a* | Gcc atc ttt caa ttg tgt tcg | Agc cat ggt aag gac acc tc |

**Analysis of protein binding to specific chromatin regions by ChIP-qPCR.**The MEFs (2-8 X $10^7$ cells for each genotype and condition) were harvested as described above for transcript analysis. Crosslinking and chromatin immunoprecipitation was performed as described (*50*). The cells were washed with PBS and crosslinked for 30 min in 2 ml ice cold 1% formaldehyde containing 2X Complete EDTA-free Protease Inhibitor Cocktail (Roche). Cross-linking was stopped by the addition of glycine to 333 mM. The MEFs were collected by 5 min centrifugation at 1,000 rpm and were washed with ice cold PBS containing 1X Complete EDTA-free Protease Inhibitor Cocktail (Roche). The crosslinked cells were flash frozen and stored at 80C.

The crosslinked cells were thawed and lysed by the addition of 4 ml ice cold cell lysis buffer (5 mM PIPES pH 8.0, 85 mM KCl, 0.5% NP-40, and 2X Complete EDTA-free Protease Inhibitor Cocktail (Roche). The nuclei were collected by 5 min centrifugation at 2,000 rpm at 4 C. The nuclei were lysed by 30 min incubation in 4 ml RIPA buffer containing 1X PBS, 1% NP-40, 0.5% sodium deoxycholate, 0.1% SDS, 20 mM N-ethylmaleimide, and 2X Complete EDTA-free Protease Inhibitor Cocktail (Roche) at 4C with rotation. The chromatin was sheared by sonicating 2 ml of the nuclear lysate in a polystyrene tube in an ice bath. All samples were sonicated the same number of cycles until the lysates became translucent (10-25 cycles, 30 s on, 30 s off, 40% duty output 2). The sonicated lysate was cleared by 14,000 rpm centrifugation at 4C for 15 min. The chromatin was divided into 0.5-1 ml aliquots (containing chromatin from 2.5 X $10^6$ – 2 X $10^7$ MEFs) for immunoprecipitation with different antibodies. The sonication procedure was optimized to produce a DNA fragment distribution with a maximum between 500 bp and 1 kb.

Each antibody (2-4 μg per reaction) and normal rabbit (Cell Signaling 2729S; Santa Cruz sc-2027) or mouse (Santa Cruz sc-2025) IgG was incubated with protein A or G Dynabeads for at least 4 h at 4C with rotation. (Invitrogen, 30-60 μl Dynabead slurry per reaction. The Dynabeads were pre-rinsed and resuspended in PBS with BSA). The beads with bound antibodies were washed and resuspended in PBS with BSA. The beads with each antibody or IgG control were added to aliquots of the cleared lysates prepared above and the samples were rotated for 24 h at 4 C. Every experiment included a negative IgG control and positive controls including H3 and transcription factors that are known to bind the regions that were tested.

The beads with bound immune complexes were washed 5 times in ice cold 100 mM Tris pH 7.5, 500 mM LiCl, 1% NP-40, 1% sodium deoxycholate, and once with ice cold 10 mM Tris-HCl pH 7.5, 0.1 mM EDTA. Chromatin complexes were eluted by incubating the beads in 1% SDS, 0.1 M NaHCO3 for 1 h at 65C and vortexing them intermittently. The beads and precipitated proteins were removed by 3 min centrifugation at 14000 rpm. The supernatant containing the eluted chromatin complexes as well as aliquots of the total input chromatin were incubated for 16-20 h at 65 C to reverse the crosslinks. The DNA was recovered using a QIAquick PCR cleanup kit (Qiagen).

The relative amounts of the promoters and flanking regions that were precipitated by each antibody were measured by qPCR. Each qPCR reaction contained 4% (vol/vol) cDNA, 1 μM of each primer, and 1X Roche SYBR Green Master Mix. The relative amounts of the regions that were analyzed were calculated by assuming that they were inversely proportional to $2^{Ct}$, where $C_t$ equals the number of cycles required to reach threshold fluorescence.

The efficiencies of precipitation (ChIP) of each region were measured relative to the amount of the region that was present in input chromatin. The amounts of the regions in the precipitated chromatin were determined by using calibration curves that were generated by analyzing serial dilutions of the corresponding input DNA samples. The average percentage of the input DNA that was precipitated (ChIP Efficiency) was plotted for duplicate qPCR reactions with upper and lower error bars corresponding to 2 times the standard deviation. The oligonucleotide primers for ChIP-qPCR were obtained from IDT, were validated by melt-curve analyses, and had the following sequences:

| **ChIP-qPCR** | **Forward primer sequence** | **Reverse primer sequence** |
|---|---|---|
| *Ifnb1* | Att cct ctg agg cag aaa gga cca | Gca aga tga ggc aaa ggc tgt caa |
| *Tnf* | Aac cct ctg ccc ccg cga tg | Tcc tcg ctg agg gag ctt ctg c |
| *Il6* | Tgg gga tgt ctg tag ctc att | Gga act gcc ttc act tac ttg c |
| *Ccng1* | Gca aaa caa aaa ccc tca cag | Cag gtc tgg cct cta tta gcc |
| *Cdkn1a* | Ttg cct ctc gga gac cag | Cca ccc agg act gaa cag a |
| *Ifnb1* -10 kb | Tcc tgc agc att cgt aca ag | Cat tcc tct ctc ccc ttg c |
| *Ifnb1* - 50 kb | Ttt ctg ttt tct gtc gga tca c | Acc aat agc gtt gag aga cca |

**Analysis of protein partitioning into nuclear and cytoplasmic fractions.** The MEFs (200,000 cells per sample) were harvested as described above for transcript analysis. The cytoplasmic and nuclear fractions were separated as described previously (*45*), with the following modifications. The MEFs were lysed by resuspension in 0.2 ml of ice cold cell lysis buffer (5 mM PIPES pH 8.0, 85 mM KCl, 0.5% NP-40, 20 mM N-ethylmaleimide, and 2X Complete EDTA-free Protease Inhibitor Cocktail (Roche)) for 30 min on ice. The nuclei were pelleted by 5 min centrifugation at 2,000 rpm at 4 C. The supernatants containing cytoplasmic proteins were collected. The nuclear pellets were washed using ice cold cell lysis buffer. The pelleted nuclei were lysed by resuspending them in 50 μl ice cold RIPA buffer (1X PBS, 1% NP-40, 0.5% sodium deoxycholate, 0.1% SDS, 20 mM N-ethylmaleimide, and 2X Complete EDTA-free Protease Inhibitor Cocktail (Roche)) and were designated nuclear extracts. The supernatant and pellet fractions were boiled in 1X SDS loading buffer 58.33 mM Tris·Cl, 5% glycerol, 1.71% SDS, 0.1 M DTT, 0.002% bromophenol blue, pH 6.8, for 10 min, centrifuged 15 min at 14,000 rpm, and equal volumes were analyzed by polyacrylamide gel electrophoresis followed by immunoblotting.

**Immunoblotting.** Fractionated cell extracts were resolved on NuPAGE 4-12% Bis-Tris protein gels (Invitrogen), transferred to PVDF membranes, and probed using the indicated antibodies. α-H3 and α-H3k9me2 were used at 1:10,000 and 1:1000 dilutions, respectively, in 1% non-fat instant milk or 5% BSA. All other antibodies were used at a dilution of 1:250. To reduce non-specific antibody binding to the membranes, they were incubated overnight at 4˚C in either 5% nonfat instant milk or 5% BSA (Fisher Scientific 50-253-916) in TBS-T [60 g Tris base, 200 g NaCl in 25 L ddH20, pH 7.6 + 0.1% Tween-20]. The membranes were incubated with primary antibody solutions overnight at 4˚C, washed 2 X 15 min with 1X TBS [60 g Tris base, 200 g NaCl in 25 L ddH20, pH 7.6] + 0.1% Tween-20 at R.T. with shaking, incubated with secondary antibody solution (goat anti-mouse or goat anti-rabbit ECL antibodies, 1:10,000) containing 1% nonfat instant milk or 1% BSA 1 h at R.T. with shaking, washed 2 X 15 min in TBS-T at R.T. with shaking, developed with SuperSignal West Femto Maximum Sensitivity Substrate (Thermo Scientific, 34095) according to the manufacturers protocol, and visualized using Bio-Rad ChemiDoc Imaging System. Background-corrected signals for each band were calculated from the recorded images using ImageJ (*51*).

**Visualization of protein complexes in living cells by using BiFC analysis.**

Plasmids were constructed that encoded the proteins of interest fused to fragments of YFP that could form BiFC complexes (*46*). The N-terminal end of Keap1 was fused to YFP residues 1-172 (YN) as well as 173-238 (YC). The C-terminal end of NFκB p50 was fused to YFP residues 1-172 (YN). The C-terminal end of NFκB p65 was fused to YFP residues 155-238 (YC). The C-terminal end of the Rel homology region of NFκB p50 (residues 42-354) was fused to YFP residues 173-238. The C- terminal end of the Rel homology region of NFκB p65 (residues 6-296) was fused to YFP residues 1-172. Plasmids that encoded the combinations of fusion proteins indicated in each experiments were co-transfected into HEK293T or MEF cells using Fugene6 according to the manufacturer's instructions. The cells were cultured on #1.5 coverglass in the same medium that was used to culture the MEFs for 24-72 h after transfection. To enhance the fluorescence intensity, the cells were cultured for 30 min to 3 h at 30°C, which facilitates maturation of the fluorescent complex. Hoechst was added to 20 μg/ml, and the cells were viewed using an Olympus IX-81 inverted microscope with a spinning disk confocal accessory at about 30°C for up to 1 h. BiFC complex fluorescence was imaged using YFP excitation and emission filters. Hoechst fluorescence was imaged using DAPI excitation and emission filters. The excitation and emission light was separated using a multi-pass dichroic mirror. Images were captured with a Hamamatsu ORCA-ER CCD camera without the use of electronic gain.

The images were acquired using Slidebook software and were analyzed using the FIJI implementation of ImageJ. Images were acquired of cells that represented the full range of different distributions that were observed in the population. In particular, cells with a wide range of different fluorescence intensities were imaged. Because the objective was to capture the full range of different distributions, it is likely that multinuclear cells and cells with BiFC signal that was associated with condensed chromatin or excluded from condensed chromatin were overrepresented in the captured images. Nevertheless, cells that expressed different BiFC complexes were subject to the same selection. The order in which different BiFC complexes were imaged was altered between different experiments. In a subset of experiments, the same sample was analyzed twice, and the results were reproducible (20% maximum change in the proportion of cells in each category). It is therefore unlikely that the procedures that were used to capture the images caused the differences in the frequencies of different subcellular distributions that were observed for different BiFC complexes.

The subcellular distribution was scored for every cell with detectable fluorescence by visual evaluation of the relative intensities of BiFC fluorescence in the area that overlapped the nuclear Hoechst fluorescence and the area immediately surrounding the nucleus. Analysis of a subset of cells after scoring indicated that cells in which the ratio of nuclear to cytoplasmic fluorescence was greater than approximately 1.2 or less than 0.7 were scored nuclear or cytoplasm enriched, respectively. Ratios between these values were scored to partition equally between the nucleus and the cytoplasm. The precise threshold ratios varied depending on the absolute fluorescence intensity, the dimensions of the cell, and the quality of the image (focus, background, uniformity of signal). Testing by persons who were blinded to the experimental design confirmed that the scoring was reproducible with differences of less than 20% in the number of cells in each category scored by different individuals.

The figures were produced by using Adobe Photoshop to convert the BiFC fluorescence intensity signal to yellow color, and the Hoechst fluorescence intensity signal to blue color. Each conversion was performed by linear transformation of the minimum signal in the region of interest to color intensity 0 and the maximum signal to color intensity 255. The images were superimposed and minor adjustment of color registration due to chromatic aberration was performed. The images were cropped to a size that accommodated the cells of interest and a margin sufficient to judge the background. The TIFF images were compressed to PDF format for presentation.

**Analysis of the effects of lysine methyltransferase inhibitors on transcription.**

BIX01294 was dissolved in filtered ddH20. The indicated concentration, typically 20 μM, of BIX01294 or vehicle was added the indicated time, typically 1 h, prior to Sendai virus infection. To analyze the effects of MS012 on transcription and protein binding, it was dissolved in DMSO and added at a concentration of 1 μM 48 h prior to Sendai virus infection. Different concentrations and times of incubation were used for the different compounds because of differences in their potencies and solubilities. No effects of MS012 in cells have been reported or observed by us prior to 48 h. When the effects of BIX01294 were compared with those of other lysine methyltransferase inhibitors, DMSO was added to the cells 48 h prior to infection.

### Materials

**Antibodies:** The αKeap1 (Aviva Systems Biology OACD04962, lot A20190909988) antibody was raised using recombinant Keap1 Met1-Gln286 for immunization. The αKeap1-N (Santa Cruz sc-15246, lot H2603) antibody has an epitope mapping near the N-terminus of human Keap1. The αKeap1-C (Proteintech 10503-2-AP, lot 00051883) antibody was raised using a recombinant human Keap1 325-624 fragment for immunization and used in (*52, 53*). The sources of the other antibodies were as follows: αp50/p105 (Cell Signaling 13586, D4P4D lot 2), αp52/p100 (Cell Signaling 4882, lot 6), αp65 (Cell Signaling 8242, lot 9), αphospho-p65-Ser536 (Cell Signaling 3031, lot 11), αG9a (Cell Signaling 3306, lot 2), αGLP (Abcam ab41969, lot GR191907-41), αH3K9me1 (Abcam ab8896, lot GR3236306-2), αH3K9me2 (Abcam ab1220, lots GR3228498-2, GR325223-3), αH3 (Abcam ab1791, lot GR3198255-1), αIκβα (Cell Signaling 4814, lot 17), αIKKβ (Cell Signaling 2678, lot 4), αphospho-IKKα/β (Cell Signaling 2697, lot 19), αphospho-TBK1 (Cell Signaling 5483, lot 8), αIRF3 (Santa Cruz sc-9082, lot G0910), αJun (Santa Cruz sc-1694, lot E0207), αvinculin (Cell Signaling 4650T, lot 4), αlaminB1 (Cell Signaling 13435, lot 2), αCul3 (Sigma C0871, lot 126M4852V), normal rabbit IgG (Cell Signaling 2729, lot 9), normal rabbit IgG (Santa Cruz sc-2027, lot L2414), normal mouse IgG (Santa Cruz sc-2025, lot I2208).

**Cell culture materials:** DMEM (Gibco 11995) was supplemented with 10% (v/v) fetal bovine serum (Atlanta Biologicals S11550 lot H1030) and 0.1 mg/ml penicillin-streptomycin (Gibco 15140). PBS (10010) and 0.05% trypsin-EDTA solution (25300) were purchased from Gibco.

**Chemicals:** The following chemicals were obtained from Cayman Chemical: BIX01294 (13124, ≥98%), GSK343 (14094 ≥95%). The following chemicals were obtained from Millipore Sigma: MS012 (SML2174 ≥98%). These compounds were dissolved in ddH2O (BIX01294) or DMSO (all other compounds) at concentrations ranging from 20 to 50 mM. The final concentration of DMSO in experiments using this vehicle was 0.1%. Buffers, salts, detergents were purchased from Millipore Sigma and Thermo Fisher Scientific.

**Statistical analyses to evaluate the likelihood of chance occurrence of observed results.** Each graph shows the means and two times the standard deviation for replicate measurements from one experiment. In addition, to determine the reproducibility of the results, data from multiple independent experiments with different MEFs of each genotype were tested by two-factor ANOVA with replication that was implemented in the Real Statistics Microsoft Excel plug-in v. 6.8 (Copyright (2013 – 2020) Charles Zaiontz. www.real-statistics.com). ANOVA analyses were performed using data that were obtained from experiments that were conducted independently, and where the mean value for one of the measurements that were compared was at least 3 fold higher than the negative control (e.g. IgG in ChIP experiments). All measurements that were compared were obtained from experiments that were conducted in parallel for the MEFs of different genotypes or for the different conditions that were compared (i.e. balanced analyses). We have not demonstrated that the measurements are normally distributed or that the variances for the two factors were the same. To compensate for potential error that could be caused by non-normal or asymmetric distributions, we rejected null hypotheses only if they had low probabilities ($p<0.001$ or lower).

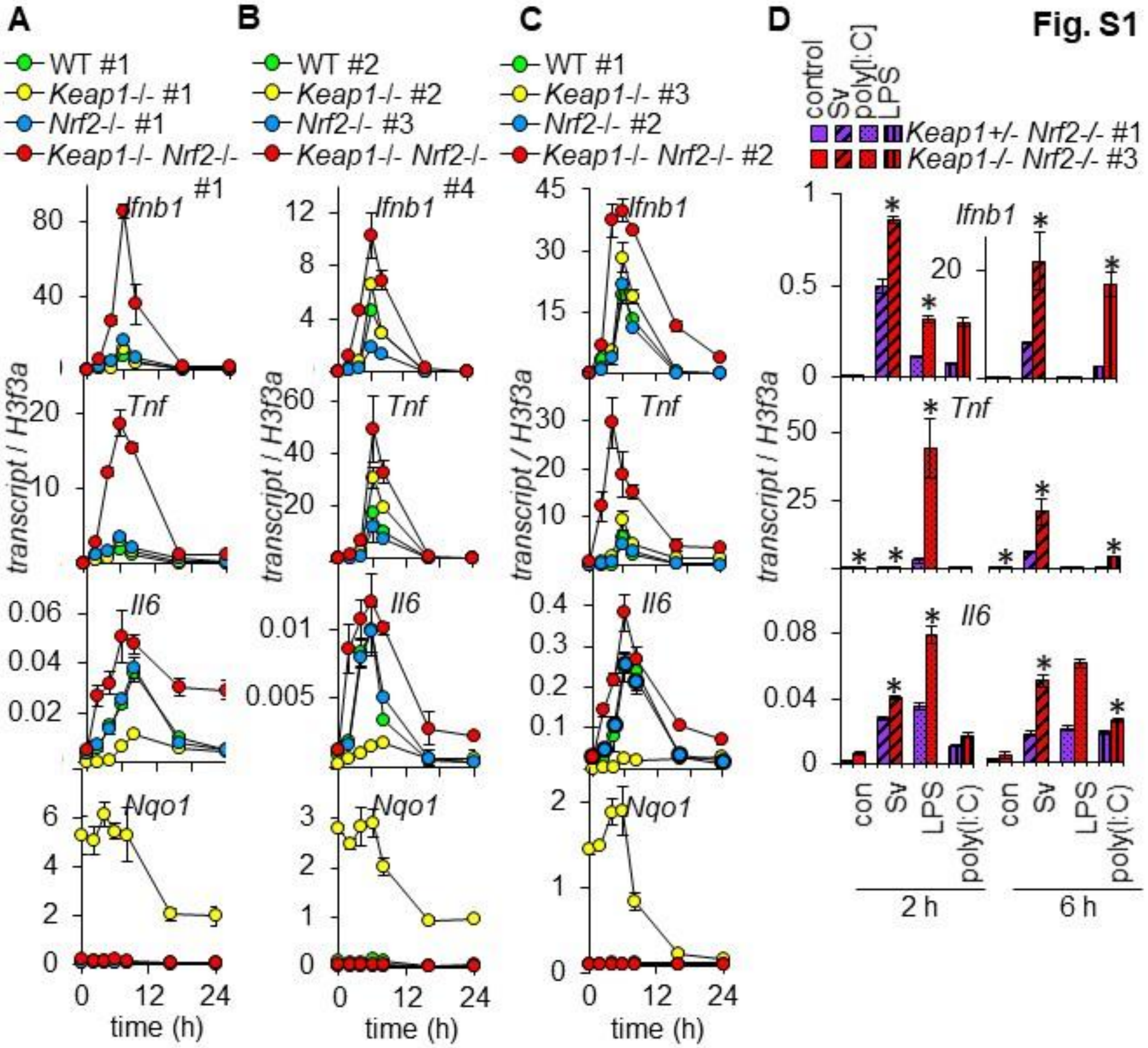

Fig. S1
A
WT #1
Keap1-/- #1
Nrf2-/- #1
Keap1-/- Nrf2-/- #1
B
WT #2
Keap1-/- #2
Nrf2-/- #3
Keap1-/- Nrf2-/- #4
C
WT #1
Keap1-/- #3
Nrf2-/- #2
Keap1-/- Nrf2-/- #2
D
control
Sv
poly[I:C]
LPS
Keap1+/- Nrf2-/- #1
Keap1-/- Nrf2-/- #3
Ifnb1
Tnf
Il6
Nqo1
transcript / H3f3a
time (h)
con
Sv
LPS
poly(I:C)
2 h
6 h

**Fig. S1. *Keap1-/-* deletions increase *Ifnb1, Tnf,* and *Il6* induction in multiple independently derived MEFs lacking Nrf2, and increase *Nqo1* transcription only in MEFs with intact Nrf2 (related to Fig. 1).**

(**A**) The levels of *Ifnb1*, *Tnf*, *Il6,* and *Nqo1* transcripts were measured at the indicated times after Sendai virus infection in wild type (WT#1), *Keap1-/-* #1, *Nrf2-/-* #1, and *Keap1-/- Nrf2-/-* #1 MEFs. The *Keap1-/-* deletions increased *Nqo1* transcription 100-fold in MEFs with intact Nrf2, whereas they had small effects on *Ifnb1*, *Tnf*, and *Il6* transcription in MEFs with intact Nrf2. In contrast, the *Keap1-/-* deletions had no significant effect on *Nqo1* transcription in *Keap1-/- Nrf2-/-* MEFs, whereas they increased virus induction of *Ifnb1*, *Tnf*, and *Il6* transcription in *Keap1-/- Nrf2-/-* MEFs. Consequently, Keap1 regulated *Nqo1 versus Ifnb1*, *Tnf*, and *Il6* transcription by distinct mechanisms. Electrophile response gene transcription did not influence the effects of Keap1 on *Ifnb1*, *Tnf*, and *Il6* transcription in *Nrf2-/-* MEFs. Part of the data are shown in Fig. 1A and are reproduced here for comparison. The experiments shown in panels A-C are representative of 7 experiments with 6 different sets of MEFs (MEFs that were isolated from different embryos are identified by #number).

(**B**) The levels of *Ifnb1*, *Tnf*, *Il6,* and *Nqo1* transcripts were measured at the indicated times after Sendai virus infection in wild type (WT#3), *Keap1-/- #3*, *Nrf2-/- #3*, and *Keap1-/- Nrf2-/- #3* MEFs and were analyzed as described in panel A.

(**C**) The levels of *Ifnb1*, *Tnf*, *Il6,* and *Nqo1* transcripts were measured at the indicated times after Sendai virus infection in wild type (WT#1), *Keap1-/- #3*, *Nrf2-/- #2*, and *Keap1-/- Nrf2-/- #2* MEFs and were analyzed as described in panel A.

(**D**) The levels of *Ifnb1*, *Tnf*, and *Il6* gene transcripts were measured in *Keap1+/- Nrf2-/-* #1 and in *Keap1-/- Nrf2-/-* #1 MEFs 2 h (left) and 6 h (right) after Sendai virus infection; after 0.2 μl/ml lipopolysaccharide addition (LPS); and after 4 μg/ml poly[I:C] addition; and in vehicle controls (con) as indicated below the bars. The experiment shown is representative of 3 experiments using 2 different pairs of MEFs.

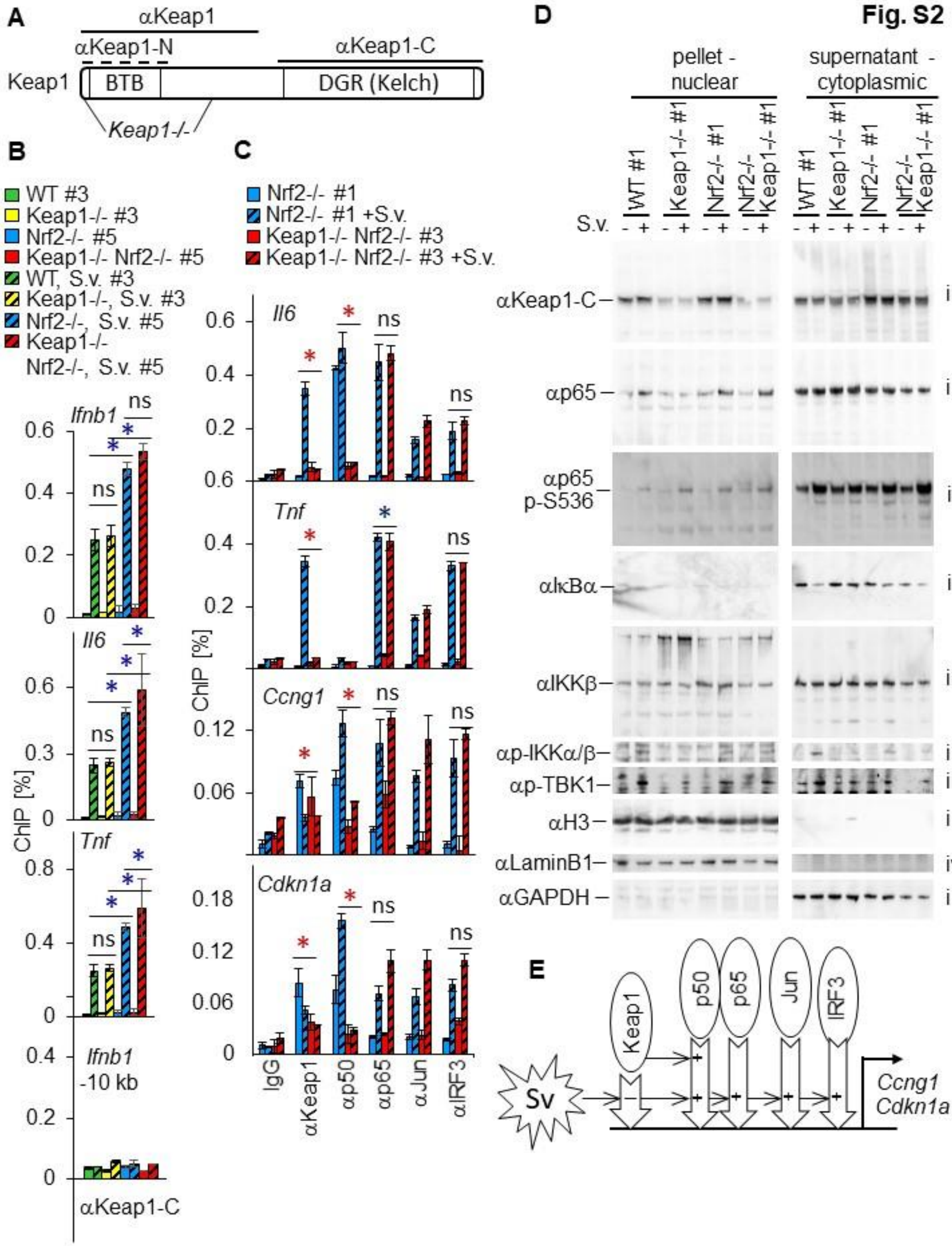

Fig. S2
A
αKeap1
αKeap1-N
αKeap1-C
Keap1
BTB
DGR (Kelch)
Keap1-/-
B
WT #3
Keap1-/- #3
Nrf2-/- #5
Keap1-/- Nrf2-/- #5
WT, S.v. #3
Keap1-/-, S.v. #3
Nrf2-/-, S.v. #5
Keap1-/- Nrf2-/-, S.v. #5
Ifnb1
Il6
Tnf
Ifnb1 -10 kb
ChIP [%]
αKeap1-C
C
Nrf2-/- #1
Nrf2-/- #1 +S.v.
Keap1-/- Nrf2-/- #3
Keap1-/- Nrf2-/- #3 +S.v.
Il6
Tnf
Ccng1
Cdkn1a
ChIP [%]
IgG
αKeap1
αp50
αp65
αJun
αIRF3
D
pellet - nuclear
supernatant - cytoplasmic
WT #1
Keap1-/- #1
Nrf2-/- #1
Nrf2-/- Keap1-/- #1
S.v.
αKeap1-C
αp65
αp65 p-S536
αIκBα
αIKKβ
αp-IKKα/β
αp-TBK1
αH3
αLaminB1
αGAPDH
E
Keap1
p50
p65
Jun
IRF3
Sv
Ccng1
Cdkn1a

**Fig. S2. Virus infection induces Keap1 binding at the *Ifnb1, Tnf,* and *Il6* promoters, but not at the *Ccng* or *Cdkn1* promoters. Keap1 is required for NFκB p50, but not for NFκB p65 recruitment, or for canonical NFκB signaling (related to Fig. 2).**

(**A**) Regions of Keap1 that were used to raise the antibodies and that were deleted by the *Keap1-/-* deletions. The dashed line indicates an epitope near the N-terminus.

The *Keap1-/-* deletions removed the sequences that encode the Keap1 regions that were used to raise the αKeap1 and αKeap1-N antibodies, but they did not remove the sequence that encodes the Keap1 region that was used to raise the αKeap1-C antibody.

(**B**) The levels of Keap1 binding at the *Ifnb1, Tnf,* and *Il6* promoters, and 10 kb upstream of the *Ifnb1* promoter, were measured in wild type (WT #3), *Keap1-/-* #3, *Nrf2-/-* #5, and *Keap1-/- Nrf2-/-* #5 MEFs after mock (solid bars) and Sendai virus (striped bars) infection using αKeap1-C antibodies. The reproducibility of the differences in the ChIP signals indicated in panels A and B were evaluated by two-factor ANOVA analyses of data from 2-9 experiments using 2-8 different sets of MEFs (* $p<0.001$ increase[blue] decrease[red], Table S1). This experiment was performed in parallel with, and used the same samples as, the experiment shown in Fig. 2A.

Several independent lines of evidence support the interpretation that the αKeap1-C antibody recognizes both Keap1 and protein(s) that are related to Keap1 in ChIP assays. First, the αKeap1-C antibody produces ChIP signals in all the regions, and only in the regions in which the αKeap1 and αKeap1-N antibodies also produce ChIP signals. Second, Nrf2 counteracts the ChIP signals that are produced by both the αKeap1-C as well as the αKeap1 and αKeap1-N antibodies, as reflected by the higher ChIP signals in *Nrf2-/-* MEFs compared to WT MEFs. Finally, BIX01294 increases the ChIP signals that are produced by the αKeap1-C as well as by the αKeap1 and αKeap1-N antibodies in *Nrf2-/-* MEFs, but not in *Keap1-/- Nrf2-/-* MEFs. Consequently, the αKeap1-C antibody recognizes Keap1 and either a Keap1 fragment or a Keap1 related protein that is (1) recruited to the same chromatin regions as Keap1, (2) whose binding to chromatin is reduced by Nrf2, and (3) whose binding to chromatin is not altered by BIX01294.

(**C**) The levels of Keap1, NFκB p50, NFκB p65, cJun, and IRF3 binding at the *Il6, Tnf, Ccng1,* and *Cdkn1a* promoters were measured in *Nrf2-/-* #1, and *Keap1-/- Nrf2-/-* #3 MEFs after mock (solid bars) and Sendai virus (striped bars) infection. This experiment was performed in parallel with, and used the same samples as, the experiment shown in Fig. 2B.

(**D**) The levels and partitioning of Keap1, NFκB p65, NFκB p65 phospho-Ser536 (p-S636), IκBα, IKKβ, phospho-IKKα/β (p-IKKα/β), phospho-TBK1 (p-TBK1), G9a, histone H3 (H3), LaminB1, and Gapdh were analyzed following hypotonic lysis of wild type (WT#1), *Keap1-/- #1*, *Nrf2-/- #1*, and *Keap1-/- Nrf2-/- #1* MEFs after mock (-) or Sendai virus (+) infection as indicated above the lanes. The pellets containing nuclear proteins (left) and the supernatants containing cytoplasmic proteins (right) were separated on parallel gels and were blotted together using the antibodies indicated to the left of the images. The same protein extracts were analyzed on several gels and membranes as indicated to the right of the images.

The level of Keap1 was higher in *Nrf2-/-* MEFs than in wild type MEFs, consistent with the higher levels of Keap1 binding to the *Ifnb1, Tnf* and *Il6* promoters in *Nrf2-/-* MEFs.

The αKeap1-C antibody detected a band that migrated with a mobility similar to Keap1 in extracts from *Keap1-/-* and *Keap1-/- Nrf2-/-* MEFs. The relationship between this band and the ChIP signals that were detected by this band in the same MEFs is not known.

(**E**) Effects of Keap1 on transcription factor recruitment to the *Ccng1 and Cdkn1a* promoters. The + symbols inside the hollow arrows indicate the factors that were required for chromatin binding by each protein. The concerted recruitment of Keap1, NFκB p50 and NFκB p65 to several promoters upon virus infection and the concordant loss of NFκB p50 binding and transcription repression in *Keap1-/- Nrf2-/-* MEFs suggest that Keap1 interactions with NFκB p50 and with NFκB p65 regulate chromatin binding and transcription.

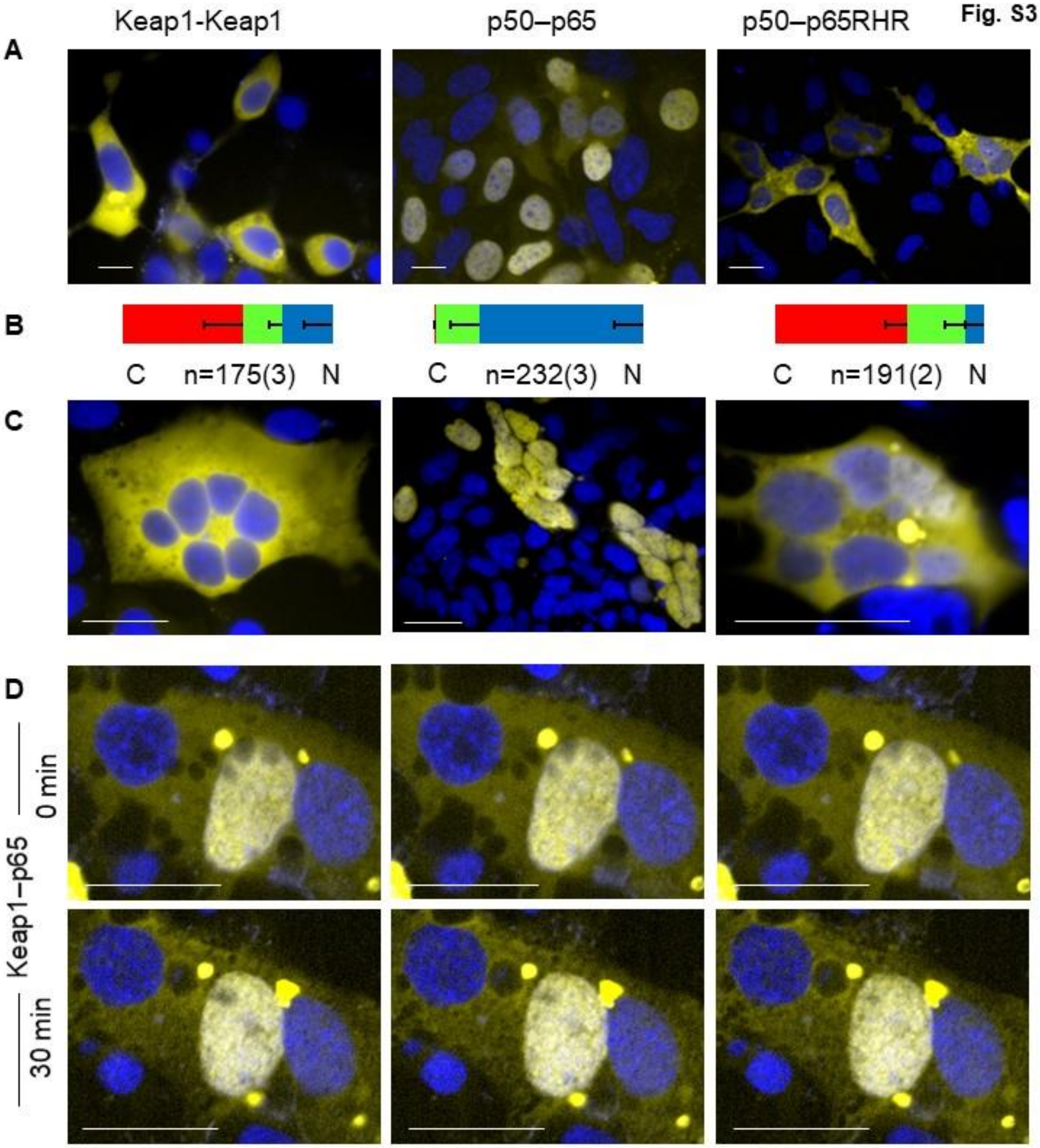
Fig. S3
Keap1-Keap1
p50–p65
p50–p65RHR
A
B
C
n=175(3)
N
C
n=232(3)
N
C
n=191(2)
N
C
D
Keap1–p65
0 min
30 min

**Fig. S3. Keap1-Keap1, p50-p65, p50-p65RHR, Keap1-p65 and Keap1-p50RHR BiFC complexes differ in subcellular distributions, in selective enrichment among individual nuclei within multinucleate cells (related to Fig. 3).**

(**A**) BiFC analyses of Keap1 homodimers, NFκB p50-p65 heterodimers, and NFκB p50 complexes with the NFκB p65 Rel homology region (RHR). Each column shows fluorescence images of cells with BiFC complexes formed by the proteins indicated at the top of the column. Each pair of proteins was fused to complementary fluorescent protein fragments, and expressed in HEK293T cells (see Materials and Methods). It is possible that the ectopic expression of fusion proteins or BiFC complex formation affects these complexes. The BiFC complex fluorescence is shown in yellow and Hoechst stain is shown in blue. The images were selected to represent the range of distributions that were observed for each BiFC complex. Scale bars 30 μm. The BiFC complexes that are shown here were analyzed in parallel with the complexes that are shown in panels 3A, 3B and 3C.

(**B**) Distributions of the BiFC complexes indicated at the top of each column. The bars show the proportions of cells (mean and sem) in which BiFC complexes were enriched in the cytoplasm (red), the nucleus (blue) and equally in the cytoplasm and the nucleus (green). n: total number of cells imaged (number of experiments in parentheses).

(**C**) Fluorescence images of multinucleate cells with BiFC complexes formed by the proteins indicated at the top of each column of images. The images were selected to represent the range of BiFC complex enrichment in different nuclei within the same multinucleate cell.

Keap1 homodimer and p50-p65 heterodimer BiFC complexes were equally depleted or enriched in all of the nuclei of multinucleate cells. This is in contrast to the differences in enrichment of Keap1-p50 and Keap-p65 BiFC complexes among different nuclei in many multinucleate cells (see Fig. 3C).

(**D**) Optical sectioning of a multinucleate cell at different times to establish the differences in the levels of Keap1-p65 BiFC complexes in different nuclei and to determine the stability of those differences over time. The cell that is shown in Fig. 3Cii was imaged at times that were separated by 30 minutes (upper and lower sets of images) using a spinning disk confocal microscope to collect optical sections that were spaced 2 μm apart (horizontal row of images). The images of BiFC (yellow) and Hoechst (blue) fluorescence were superimposed.

**Fig. S4**

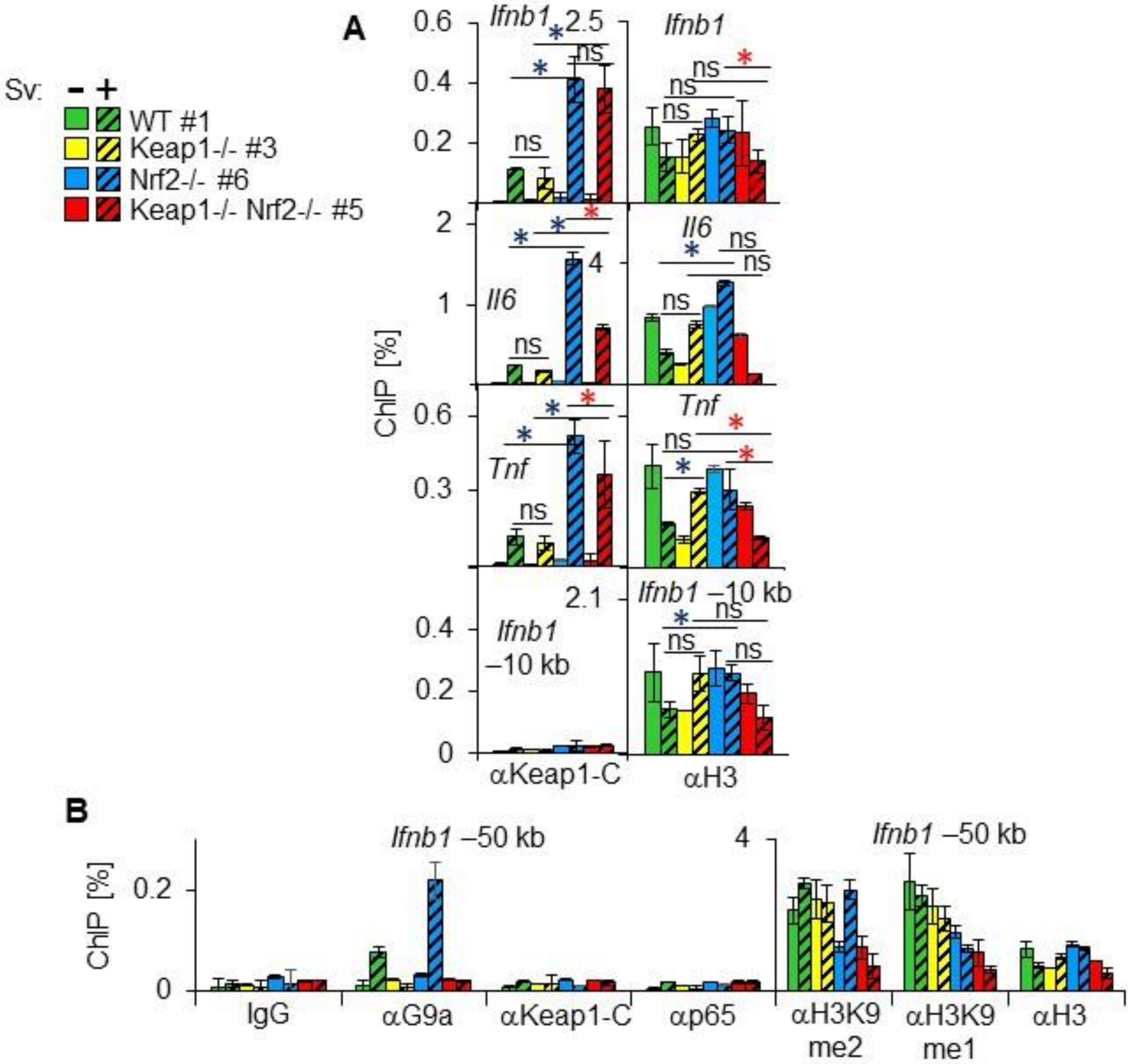
A
B
Sv: – +
WT #1
Keap1-/- #3
Nrf2-/- #6
Keap1-/- Nrf2-/- #5
ChIP [%]
*Ifnb1*
*Il6*
*Tnf*
*Ifnb1* –10 kb
ns
αKeap1-C
αH3
*Ifnb1* –50 kb
IgG
αG9a
αp65
αH3K9 me2
αH3K9 me1

**Fig. S4. Keap1 is required for G9a recruitment and for H3K9 methylation throughout the *Ifnb1* locus upon virus infection (related to Fig. 4).**

(**A**) The levels of Keap1 (αKeap1-C) and H3 binding at the *Ifnb1*, *Tnf*, and *Il6* promoters were measured in wild type (WT#1), *Keap1-/-* #3, *Nrf2-/-* #6, and *Keap1-/- Nrf2-/-* #5 MEFs 9 h after mock (solid bars) and Sendai virus (striped bars) infection. A region 10 kb upstream of the *Ifnb1* promoter was analyzed in parallel (bottom graphs). The reproducibility of the differences in the ChIP signals indicated in panels A and B were evaluated by two-factor ANOVA analyses of data from 2-9 experiments using 2-8 sets of MEFs (* $p<0.001$ increase[blue] decrease[red], Table S1). This experiment was performed in parallel with, and used the same samples as, the experiment shown in Fig. 4A.

(**B**) The levels of Keap1, G9a, and p65 binding, and of H3K9me2, H3K9me1, and total H3 in a region 50 kb downstream of the *Ifnb1* promoter were measured in wild type (WT#1), *Keap1-/-* #3, *Nrf2-/-* #6, and *Keap1-/- Nrf2-/-* #5 MEFs 9 h after mock (solid bars), and Sendai virus (striped bars) infection. This experiment was performed in parallel with, and used the same samples as, the experiment shown in Fig. 4A.

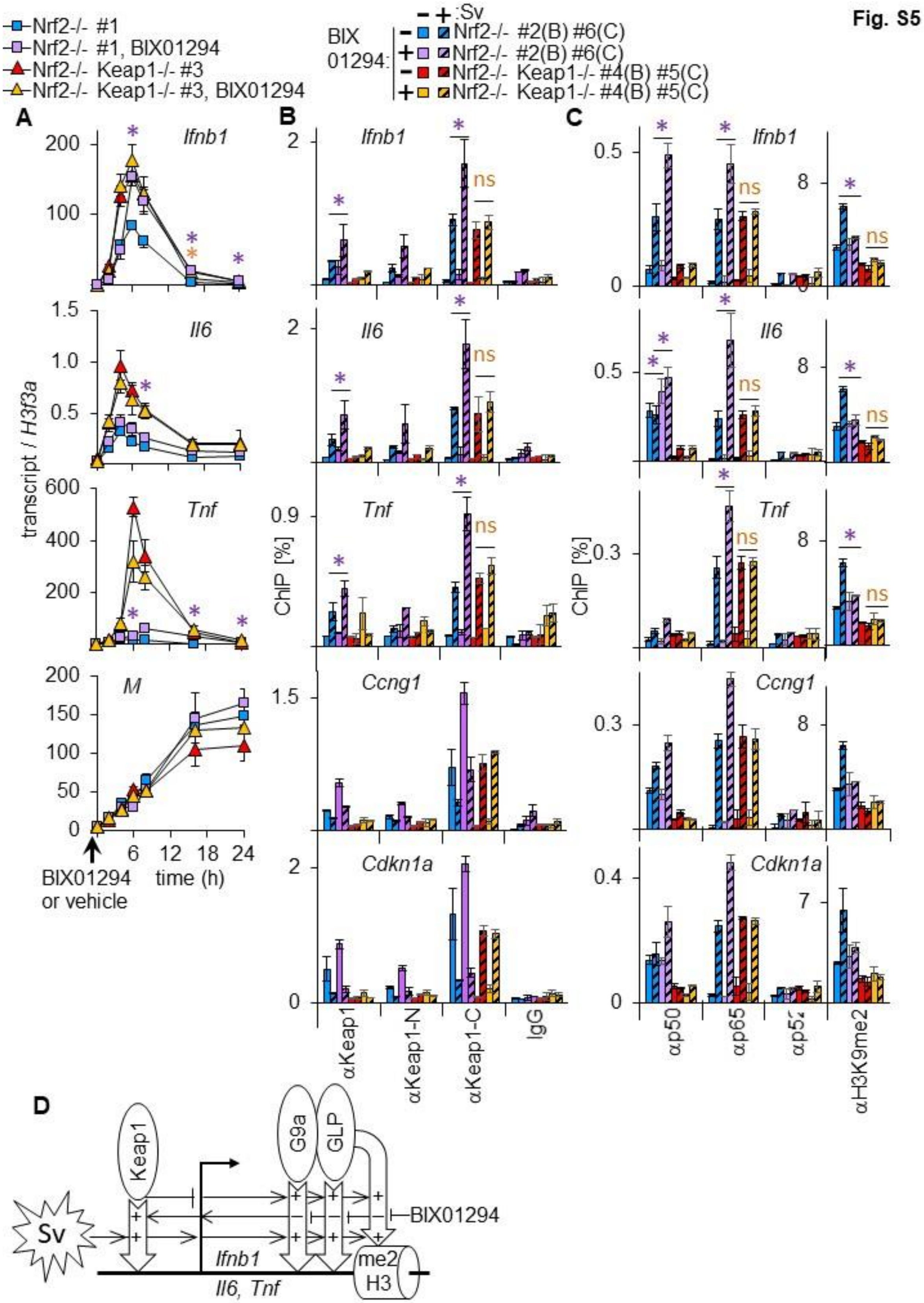

Fig. S5
Nrf2-/- #1
Nrf2-/- #1, BIX01294
Nrf2-/- Keap1-/- #3
Nrf2-/- Keap1-/- #3, BIX01294
BIX 01294
:Sv
Nrf2-/- #2(B) #6(C)
Nrf2-/- Keap1-/- #4(B) #5(C)
A
B
C
D
Ifnb1
Il6
Tnf
M
Ccng1
Cdkn1a
transcript / H3f3a
ChIP [%]
BIX01294 or vehicle
time (h)
ns
αKeap1
αKeap1-N
αKeap1-C
IgG
αp50
αp65
αH3K9me2
Keap1
G9a
GLP
Sv
BIX01294
Ifnb1
Il6, Tnf
me2
H3

**Fig. S5. BIX01294 enhances virus induction of transcription and Keap1 and NFκB binding at cytokine promoters selectively in MEFs with intact Keap1 (related to Fig. 5).**

(**A**) The levels of *Ifnb1, Il6, Tnf* and *M* gene transcripts were measured in *Nrf2-/-* #1, and in *Keap1-/- Nrf2-/-* #3 MEFs at the indicated times after Sendai virus infection following 20 μM BIX01294 or vehicle addition 1 h before infection (indicated by arrowhead on the horizontal axis). The reproducibility of the effects of BIX01294 on transcription were tested by two-factor ANOVA analyses of data from 2 experiments with different pairs of MEFs (* $p<0.0001$; purple symbols refer to *Nrf2-/-* MEFs, orange symbols refer to *Keap1-/- Nrf2-/-* #3 MEFs; Table S1).
*Keap1-/-* deletions and BIX01294 both enhanced *Ifnb1, Il6,* and *Tnf* induction by virus infection, but they did not have an additive effect. These results suggest that *Keap1-/-* deletions and BIX01294 enhanced transcription by overlapping mechanisms.

(**B**) The levels of Keap1 binding at the *Ifnb1, Il6, Tnf, Ccng1,* and *Cdkn1a* promoters, were measured in *Nrf2-/-* #2, and *Keap1-/- Nrf2-/-* #4 MEFs 6 h after mock (solid bars) and Sendai virus (striped bars) infection following 20 μM BIX01294 or vehicle addition one hour before infection. Keap1 binding was analyzed using the αKeap1, αKeap1-N and αKeap1-C ChIP antibodies. This experiment was performed in parallel with and used the same samples as the experiment shown in Fig. 5C. The reproducibility of the effects of BIX01294 on the ChIP signals indicated in panels B and C were evaluated by two-factor ANOVA analyses of data from 3 experiments using different pairs of MEFs (* $p<0.0005$ *Nrf2-/-* [purple] *Keap1-/- Nrf2-/-* [orange], Table S1).

(**C**) The levels of NFκB p50, NFκB p65, and NFκB p52 binding and of H3K9me2 at the *Ifnb1, Il6, Tnf, Ccng1, and Cdkn1a* promoters were measured in *Nrf2-/-* #6, and *Keap1-/- Nrf2-/-* #5 MEFs 6 h after mock (solid bars) and Sendai virus (striped bars) infection following 20 μM BIX01294 or vehicle addition one hour before infection. The data are plotted in 2 parallel graphs with scales plotted to the left of each set of data. This experiment was performed in parallel with and used the same samples as the experiment shown in Fig. 5B.

(**D**) Reciprocal effects of Keap1 and of BIX01294 on G9a-GLP binding, H3K9me2, and transcription at the *Ifnb1*, *Il6* and *Tnf* genes.

Fig. 6

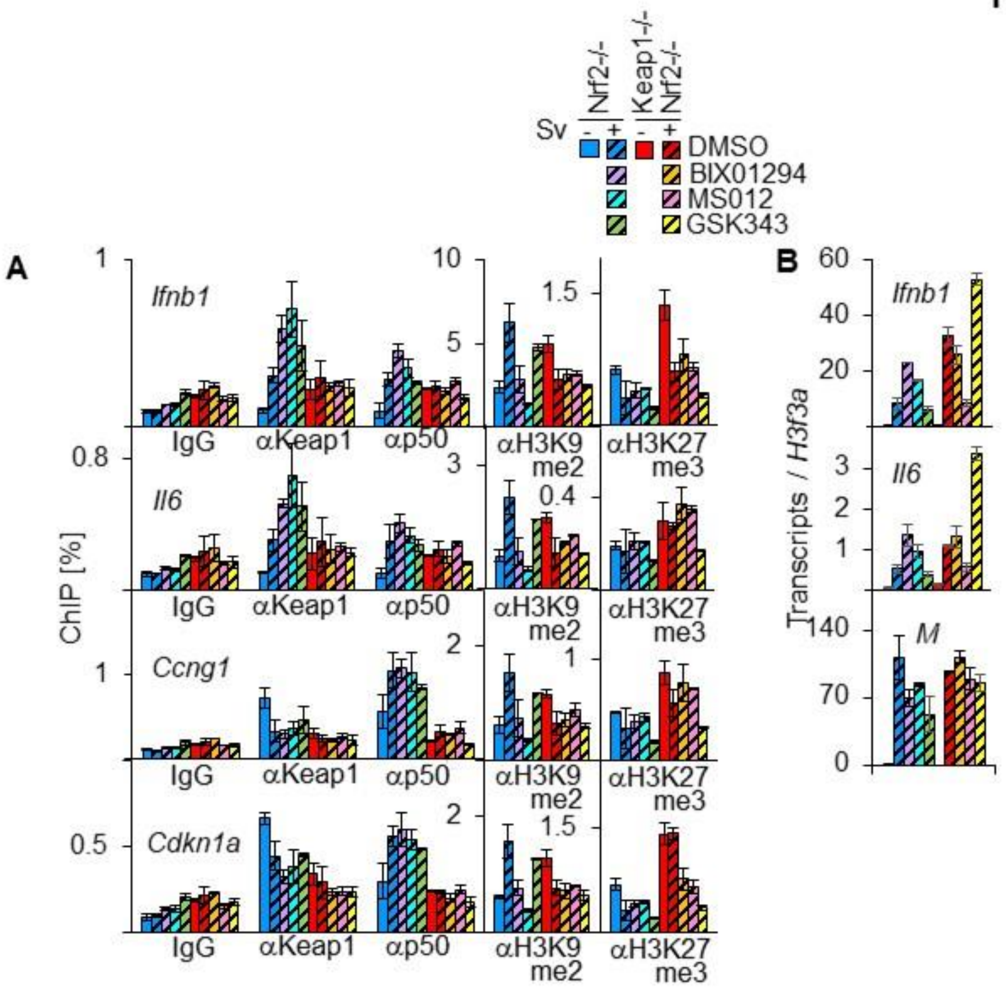

Sv
Nrf2-/-
Keap1-/-
Nrf2-/-
-
+
-
+
DMSO
BIX01294
MS012
GSK343
A
B
ChIP [%]
Ifnb1
Il6
Ccng1
Cdkn1a
IgG
αKeap1
αp50
αH3K9 me2
αH3K27 me3
Transcripts / H3f3a
M

**Fig. S6. G9a-GLP lysine methyltransferase inhibitors enhance Keap1 and NFκB p50 binding in *Nrf2-/-* MEFs selectively.**

(**A**) The levels of Keap1 and NFκB p50 binding, and H3K9me2 and H3K27me3 at the *Ifnb1, Il6, Ccng and Cdkn1a* promoters were measured in *Nrf2-/-* and *Keap1-/- Nrf2-/-* MEFs 6 hours after mock (solid bars) and Sendai virus (striped bars) infection following 1 h culture with 20 μM BIX01294, or 48 h culture with 1 μM MS012.

(B) The levels of Ifnb1, Il6, and M transcripts were measured in Nrf2-/- and in Keap1-/- Nrf2-/- MEFs that were cultured in parallel with the MEFs that were analyzed in panel A. Each graph in this Fig. shows the means and the standard errors of duplicate samples from one experiment.
